# A Bag-Of-Motif Model Captures Cell States at Distal Regulatory Sequences

**DOI:** 10.1101/2024.01.03.574012

**Authors:** Paola Cornejo-Páramo, Xuan Zhang, Lithin Louis, Yi-Hua Yang, Zelun Li, David Humphreys, Emily S. Wong

## Abstract

Deciphering the intricate regulatory code governing cell-type-specific gene expression is a fundamental goal in genetics. Current methods struggle to capture the complex interplay between gene distal regulatory sequences and cell context. We developed a computational approach, BOM (Bag-of-Motifs), which represents cis-regulatory sequences by the type and number of TF binding motifs it contains, irrespective of motif order, orientation, and spacing. This simple yet powerful representation allows BOM to efficiently capture the complexity of cell-type-specific information encoded within these sequences. We apply BOM to mouse, human, and zebrafish distal regulatory regions, demonstrating remarkable accuracy. Notably, the method outperforms more complex deep learning models at the same task using fewer parameters. BOM can also uncover cross-species sequence similarities unrecognized by genome alignments. We experimentally validate our *in silico* predictions using enhancer reporter assay, showing that motifs with the most significant explanatory power are sequence determinants of cell-type specific enhancer activity. BOM offers a novel systematic framework for studying cell-type or condition-specific cis-regulatory sequences. Using BOM, we demonstrate the existence of a highly predictive sequence code at distal regulatory regions in mammals driven by TF binding motifs.

## Introduction

Understanding the mechanisms by which genetic information is converted into phenotype is one of the fundamental goals of biology. Central to this process is the intricate regulatory network that governs gene expression, ensuring that the correct genes are expressed at the right time and place and in the appropriate amount. Cis-regulatory elements (CREs) are regulatory DNA sequences that include enhancers, promoters, insulators, and silencers. Most disease-associated genetic variants are located at non-coding regions and are enriched at CREs ^1^.

A significant proportion of CREs act as enhancers. Enhancers are integrators of spatiotemporal information and function in determining lineage and maintaining cell identity by modulating gene expression ^2–4^. They are typically located distally to the genes whose expression they modulate. Enhancers recruit transcription factors (TFs) to their DNA sequence. These binding sites often correspond to short sequences known as ‘motifs’ (∼8-10 bp in length), which are recognized by the DNA-binding protein domains. TF binding sites are considered the minimal unit for cis-regulatory activity ^5^. TF binding motif numbers can predict the strength of TF binding ^6^.

Understanding the complex regulatory code governing cell-type-specific gene expression remains a central challenge, as the relationship between distal regulatory sequence and cell context is difficult to capture. Examining statistically overrepresented DNA recognition motifs using position weight matrices (PWMs) ^7–9^ has proven helpful for identifying regulatory elements and predicting their relationship to gene expression in invertebrate genomes, including in placozoans and poriferans ^10^, *Drosophila* ^11^ or *C.elegans* ^12^. However, due to their regulatory complexity, vertebrate enhancers have proved much more challenging. This can be attributed to the vast landscape of potential regulatory elements in large genomes, coupled with the expansion of TF gene families leading to greater gene regulatory network complexity ^13^. As a result, TF binding motifs are too degenerate and numerous throughout the genome to help predict enhancers ^14^. In mammals, 10-30% of the genome is annotated by enhancer-associated histone marks based on integrative histone mark analyses ^15^.

Advancements in genome-scale experimental profiling of the regulatory genome have enabled researchers to map CREs across diverse mammalian cell types systematically. This progress has been made possible by genome-wide techniques such as ATAC-seq (assay for transposase-accessible chromatin with sequencing) and ChIP-Seq (chromatin immunoprecipitation with sequencing) of TF binding sites and histone marks that are indicative of enhancer activity. Comparative analyses of these genome-wide datasets have revealed that cell-type specific TF binding sites and candidate enhancer regions evolve rapidly across species ^16–18^. While conserved sequences may encode functional constraints, constrained regulatory functions can be encoded in sequences showing little sequence conservation ^14,19,20^. Hence, sequence conservation, a powerful strategy for identifying coding sequences across species, is insufficient for annotating many putative enhancers^16,17,21–23^.

In recent years, deep learning models have been used to learn regulatory patterns in genomic DNA ^15,24,25^. Due to their ability to learn complex patterns from large datasets, deep learning models trained on large-scale genome-wide experiments, e.g., thousands of human and mouse epigenetic and transcriptional profiles across hundreds of cell types/lines and time points ^15^, have been used to infer regulatory signals from DNA sequences ^26–32^. Applications range from predicting the impact of variant perturbations on TF binding, chromatin state, and chromatin accessibility (e.g., DeepSea ^33^, Sei ^30^, Basenji ^34^, BPNet ^32^) to the sequence vocabulary of transcriptional initiation sites ^27^. Recent methods have also focused on inferring how changes in the genomic sequence alter gene expression (e.g., Enformer ^26^, Borzoi ^31^). K-mer-based feature enumeration has been a popular method for sequence classification and variant effect prediction by combining support vector machines (SVMs) with gapped k-mer kernel functions in gkmSVM and LS-GKM ^35–40^.

Here, we show that a sequence-based vocabulary of distal regulatory activity can be learned using an enumerative method based on PWMs by leveraging gradient-boosted trees. BOM represents distal regulatory sequences by the type and number of TFBS they contain, irrespective of order, orientation, or spacing. This approach outperforms other methods, including deep learning models. BOM reveals the existence of a highly predictive sequence code at vertebrate distal regulatory regions.

## Results

### BOM uses gradient-boosted trees to predict the cell-state-specific activity of cis-regulatory elements

We propose that the flexible bag-of-motif (BOM) approach, representing each regulatory element as a set of motif counts, can capture the sequences associated with cell-type-specific regulatory activity in a highly predictive manner ^20,41–45^. Based on this concept, we develop a computational approach called BOM to discover and interpret cell-type specific cis-regulatory sequences. The approach classifies cis-regulatory sequences based on TF motif composition. At a minimum, BOM requires two conditions for classification. These conditions can be cell states defined based on differential chromatin accessibility or histone mark assays in bulk or single cells. It uses the gradient boosting algorithm, XGBoost (Extreme Gradient Boosting) ^46^ for classification or regression, and SHAP scores to interpret how each motif contributes to the prediction.

We applied BOM to classify candidate CREs among 17 annotated cell types in mouse embryos at developmental stage E8.25 using single nucleus ATAC-seq data (snATAC-seq) ^47^. We defined CREs as distal (>1 kb from TSS) and non-exonic open chromatin regions. ATAC-seq peaks are reliable predictors of *in vivo* TF binding ^48^. Based on the strength of the regulatory signal, CREs were grouped based on cell-type specific activity, resulting in 12,079 non-overlapping sequences of length 500 bp (**Methods and Supplementary Table 1**). Motifs were annotated using GimmeMotifs ^49,50^, a motif database where motifs are clustered to reduce redundancies (**Methods**). A motif count matrix was used to train the classifier, where each sequence was represented as an unordered list of motifs (‘bag’). The frequency of each motif was recorded in an input matrix (**Fig. 1a** **and Supplementary Fig. 1**). On average, ∼89% of CREs in our analyses were annotated by motifs.

**Fig. 1.**
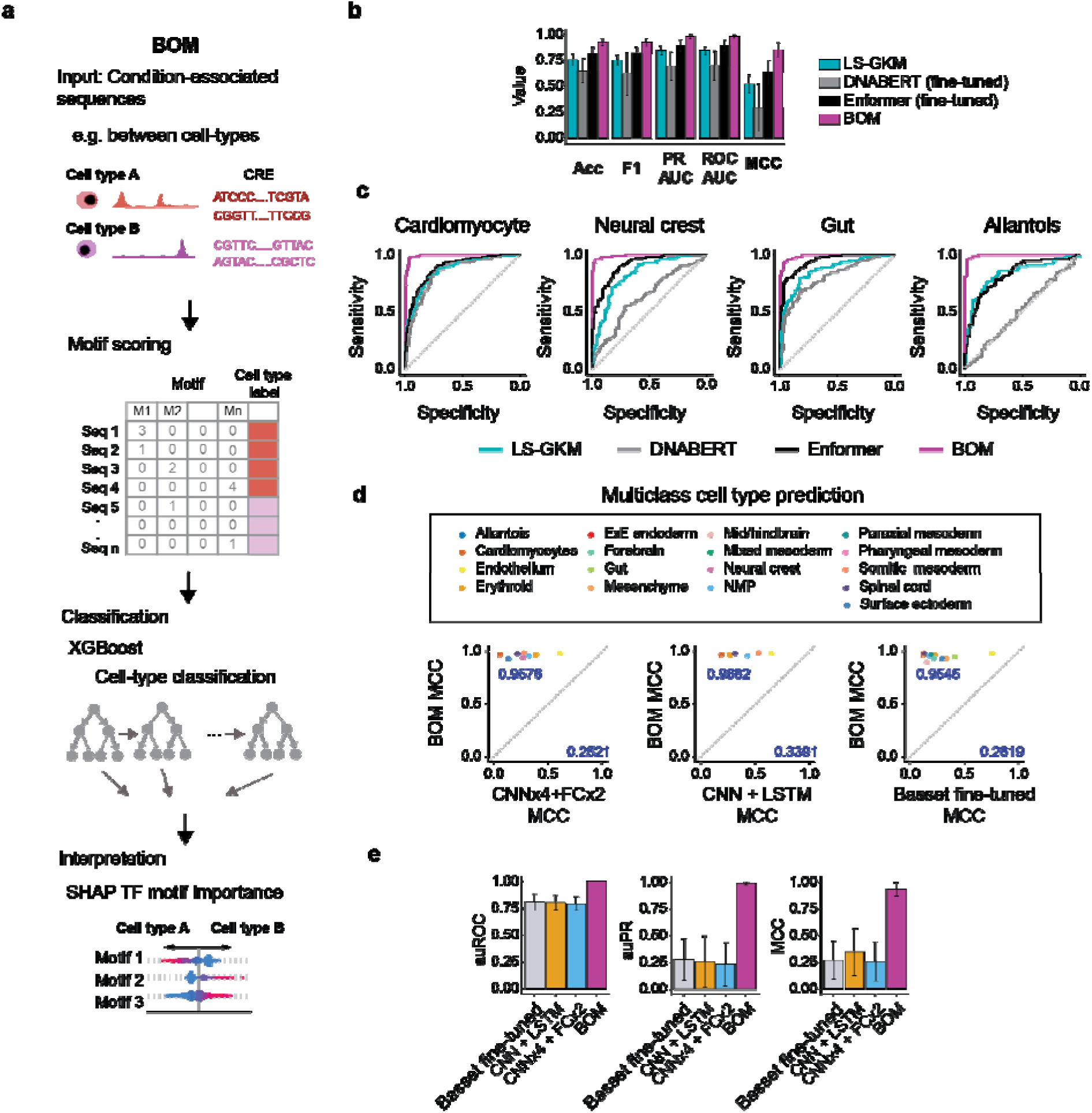
BOM overview and methods comparison. **a.** Framework of BOM (bag-of-motifs). Firstly, we define at least two sets of CRE sequences specific to different cell types or conditions. Then, we identify TF binding motif instances within these sequences. Vertebrate motifs from GimmeMotifs were used to annotate CREs. The model is trained to classify cell states using XGBoost for binary or multiclass classification. SHAP values are calculated to explain the importance of TF binding motifs for the classification task. **b**. Comparison of BOM binary classifiers using DNABERT ^52^, Enformer ^26^ and LS-GKM ^36^ across 16 mouse embryonic cell-type specific scATAC-seq peaks ^47^. Enformer and DNABERT were fined-tuned (**Methods**). Error bars represent standard deviation. **c.** ROC curves for four (of the 16) mouse cell types. **d**. Multiclass classification between mouse embryonic cell-type specific scATAC-seq peaks ^47^. Matthews correlation coefficient (MCC) for BOM versus the fine-tuned Basset model (CNNx3+FCx2) ^28^, two deep learning architectures trained from scratch, namely CNNx4+FCx2 (e.g., DeepSTARR ^53^), CNN+LSTM (e.g., DeepMEL ^54^, DanQ ^55^). **e**. Summary of model performance for models in **d**. Error bars show standard deviations.

We assessed model performance by evaluating the ability of the trained model to predict unseen test data. For binary classification, the background set of cell-type specific CREs was constructed by sampling CREs such that each cell type was equally represented in the background set. We balanced the number of CREs from the positive and negative (background) classes.

BOM performed highly in classifying 2,489 test sequences across 17 cell types by accurately classifying CREs to their respective cell types in 93% of the held-out sequences. BOM showed an average precision of 93%, recall of 92%, and an F1 score of 92% across cell types (auROC=0.98, auPR=0.98) (**Supplementary Table 2**). It also demonstrated high consistency across five random training samples, validation, and test sets (**Supplementary Fig. 2a and Supplementary Table 3**). In multiclass classification, BOM obtained an average precision of 0.99, recall of 0.88, F1 of 0.93, and auPR of 0.99 across the 17 cell types (**Supplementary Table 4**).

As cells during embryonic development can also be viewed as a continuum of transient intermediate states, we further applied BOM to understand chromatin context in intermediate cell states within the E8.25 data by using BOM to classify across 93 latent cell states in a multiclass classification ^51^ (n=76,607 CREs, mean auPR = 0.86, mean F1 = 0.71, **Supplementary Fig. 2b-c, Methods,** and **Supplementary Table 5**).

### BOM outperforms other methods in identifying the cell-type-specific cis-regulatory code

We evaluated the performance of BOM against other methods, LS-GKM, DNABERT, and Enformer, on the same distal regulatory region classification task. LS-GKM ^36^ is a gapped *k*-mer-based support vector machine approach, DNABERT ^52^ is a k-mer-based transformer-based approach, and Enformer is a large deep learning sequence model trained on over 7,000 regulatory experiments that uses dilated convolutional layers and the concept of attention to capture long-range genomic relationships ^26^. To compare the methods, we performed binary classifications between the 17 mouse developmental cell types (**Fig. 1** **b-c and Supplementary Fig. 2d**). As before, we created a balanced stratified negative set for model training. DNABERT and Enformer were fine-tuned, a process where a pre-trained model is retrained on a new dataset, which can improve its performance by leveraging the underlying pre-trained weights (**Methods**). Hyperparameter searching was performed for all neural networks. On average, BOM achieved high performance across all cell types (PRAUC=0.988, MCC=0.927). BOM outperformed LS-GKM, DNABERT, and Enformer in precision-recall AUC by 17.2%, 55.1%, and 10.3%, and in Matthews correlation coefficient (MCC) by 77.5%, 211.9%, and 33.4%, respectively (**Fig. 1b-c, Supplementary Table 2**).

Next, we compared BOM’s performance for multiclass classification to three convolutional neural network (CNN)-based deep learning architectures. These encompassed pure CNN architecture ^28,53^ and hybrid architectures that combined CNNs with recurrent neural networks (RNNs) ^54,55^; including DeepMEL & DanQ (CNN + LSTM) ^54,55^, DeepSTARR (CNNx4 + FCx2) ^53^, and Basset (CNNx3 + FCx2) ^28^.

We implemented these architectures and trained models using the mouse development data ^47^. To adapt to our task, we modified the output layer and set the number of units to 17, allowing the model to predict between the 17 cell types. We augmented the data by including their reverse complement sequences to reduce data sparsity and improve performance. Along with training from scratch, we also fine-tuned Basset and DeepMel to leverage their trained weights; we did not fine-tune DeepSTARR due to the evolutionary divergence between the original data (insects) and mammals. In each case, we used the most performant model for comparison to BOM. A hyperparameter search was done for each model.

BOM significantly outperformed the three CNN-based deep learning model architectures CNNx3 + FCx2, CNN + LSTM, and CNNx4 + FCx2, including the fine-tuned Basset model in a multiclass classification to learn cell-type specificity of distal regulatory elements (**Fig. 1d-e, Supplementary Fig. 3**). In particular, the deep learning models performed poorly on recall (range of 0.0-0.5), while BOM showed high precision and recall (average 0.98 and 0.88, respectively) (**Fig. 1d** and **Supplementary Table 6-7)**.

We also evaluated the ability of BOM models to perform well on unseen data from an independent dataset. We used mouse E8.25 BOM models to predict enhancers in a snATAC-seq dataset from a closely related developmental stage, stage E8.5 ^56^ (**Supplementary Fig. 2e**). After removing 19 cell types that were not present or annotated in the E8.25 dataset, we constructed binary classification models for the remaining 15 matched cell types from E8.5. The models achieved a mean auPR of 0.85, mean auROC of 0.76, and mean F1 score of 0.79, demonstrating that they can generalize well despite differences in cell type composition and morphology between the two embryonic times (**Supplementary Table 8**).

### Lenient motif detection threshold improves predictive performance

Determining the appropriate thresholds for motif discovery using PWMs is a recognized challenge ^9,57^. We tested different thresholds to detect motifs. Using three different thresholds for motif detection with FIMO ^50^, q-value <= 0.1, <= 0.3, and <= 0.5, we found a stricter q-value cut-off yielded decreased predictive ability in our models trained with human cell lines data (**Supplementary** Fig. 4 **and Supplementary Table 9**). With a q-value threshold of 0.5, ∼83% of the total base pairs within each distal element were annotated as motifs. Thus, most bases within CREs contain information that can be used for prediction.

This finding was corroborated by our analysis of TF binding signals using bioChIP-seq data of the corresponding TFs for six cardiac developmental TFs (Mef2a, Mef2c, Nkx2-5, Srf, Tbx5, Tead) ^58^. By calculating the mean TF binding signal across each TF’s ChIP-seq summits and matching identified motifs to corresponding TFs, we observed that a q-value threshold greater than 0.1 was necessary to capture any binding sites for Nkx2.5 and Tead1 (**Supplementary Fig. 5**). Consistent with this, prediction performance, particularly model sensitivity, was reduced by over 50% if overall motif numbers were reduced by only 10% via subsampling (**Supplementary Fig. 6**). It is possible that a less stringent threshold may enable the model to incorporate degenerate and lower-affinity motifs ^7,68^, including secondary and non-canonical motifs ^6^. However, the optimal cut-off will depend on the number and length of input data.

We further investigated the impact of eliminating overlapping motif annotations on model performance. To filter the motifs, we ordered them based on their start positions along the sequences. Next, we systematically removed overlapping motifs with lower match scores until no motifs overlapped. When two overlapping motifs had equally highest scores, one was selected at random. We observed a noticeable decrease in the prediction ability of the classifier if overlaps were removed (difference in auROC = 0.32, **Supplementary** Fig. 7 **and Supplementary Table 10**). Overlapping motifs may augment the algorithm’s ability to recognize composite arrangements of sequence motifs, as binding sites among cooperatively binding TF pairs can form composite sites that differ from the recognition sites of individual motifs ^59^. Model performance was generally sensitive to a reduction in motif counts (**Supplementary Fig. 6**).

### Distal regulatory motifs distinguish cell context across species

Beyond mouse data, we applied the Bag-of-Motifs (BOM) approach to classify cis-regulatory elements across diverse biological contexts and species (**Methods, Supplementary Table 1**). All models were trained using classed-balanced data with a stratified negative set so that cell states with low CRE numbers were not underrepresented.

We used BOM to classify 167,148 enhancer sequences between six human ENCODE cell lines with high performance (mean F1: 0.92, auPR: 0.98, auROC: 0.98; **Fig. 2a** **and Supplementary Table 11**) ^60^. Here, CREs were defined by enhancer-associated histone marks as defined using ChromHMM.

**Fig. 2.**
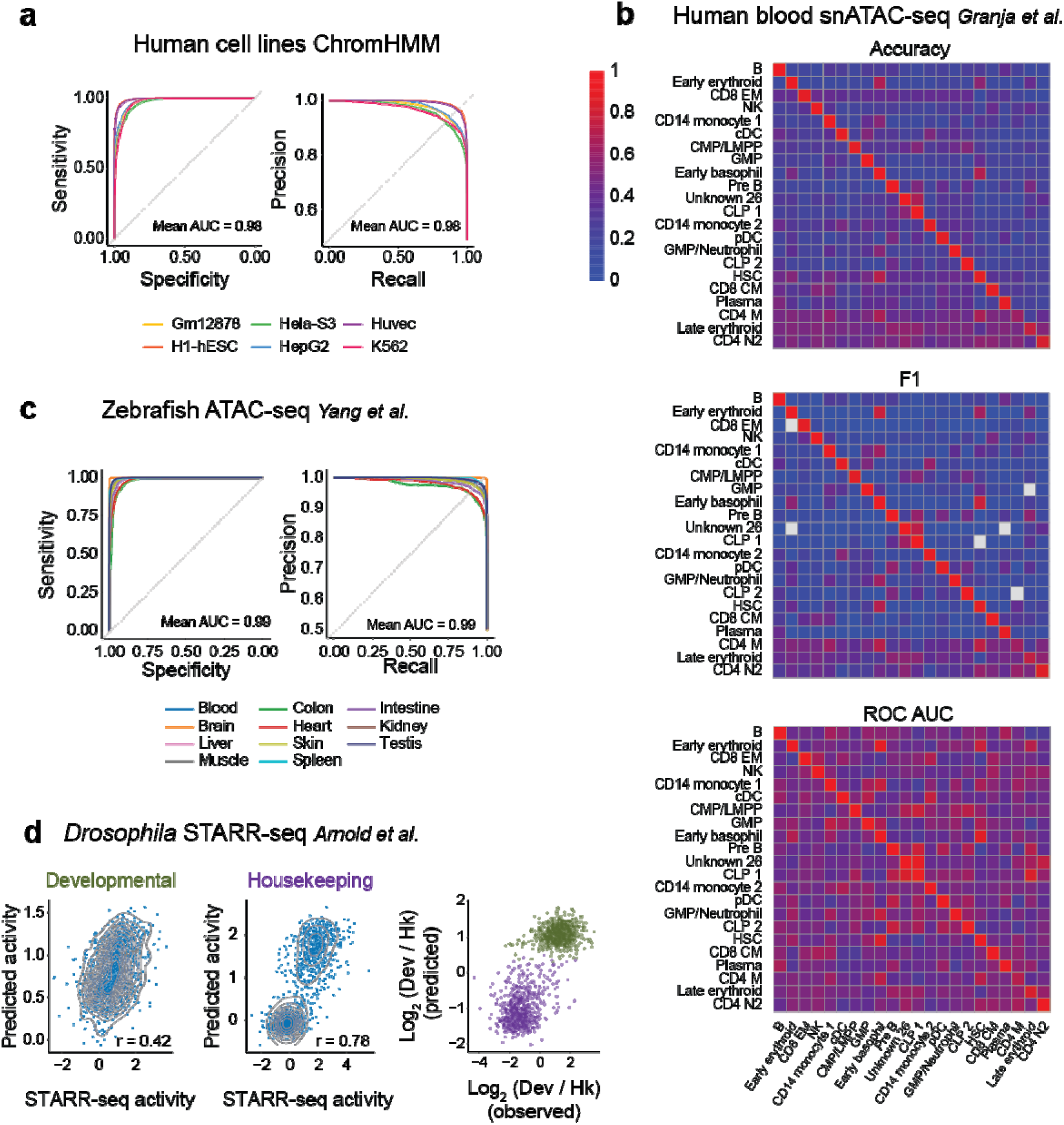
BOM accurately classifies context-specific CREs in different datasets. **a**. ROC curves (left) and precision-recall curves (right) illustrating the performance of binary BOM models in predicting cell line-specific candidate enhancers across six human cell lines (Gm12878, H1-hESC, Hela-S3, HepG2, Huvec, K562) (n = 66863). The cell line-specific candidate enhancer regions were defined using a 25-state ChromHMM model ^60^. **b**. Performance evaluation of binary BOM models trained to distinguish cell type-specific CREs for 22 human blood and bone marrow cell types ^92^. Models were trained to distinguish cell type-specific CREs from a background of CREs specific to other cell types. We evaluated the models ability to predict CREs from non-target cell types. This was done by scoring the CREs in balanced datasets CREsated for other binary models. To ensure unbiased analysis, CREs that were part of the training or validation sets of the model were excluded from the cross-dataset analysis. Accuracy, F1 scores, and auROC values were computed for each binary model (rows) and dataset (columns) (n = 5124). **c**. ROC curves (left) and precision-recall curves (right) show the prediction of tissue-specific CREs defined for 11 adult zebrafish tissues using bulk ATAC-seq data ^94^. A binary BOM model was trained for every tissue. The tissues include blood, brain, liver, muscle, colon, heart, skin, spleen, intestine, kidney, and testis (n = 59553). **d.** Correlation between mean developmental (Dev) and housekeeping (Hk) enhancer activity, as measured by MPRA in fruit fly S2 cells, and predicted activity (left and mid panels) (n = 1 258, 1 258; Dev and Hk enhancers) ^53,61^. The log_2_ fold change of Dev versus Hk enhancers for the measured activities on MPRA and the predicted values (right panel). Enhancers colored based on observed class.

We then used BOM to classify 12,796 CREs identified using snATAC human blood and bone marrow data across 22 cell types. With class-balanced data, BOM achieved a mean F1 score of 0.90, auPR of 0.95, and auROC of 0.96 (**Fig. 2b**). By testing each cell-type specific model against the regulatory elements from the other cell types, we confirmed that the models had learned a specific cis-regulatory code for each cell type (**Fig. 2b** and **Supplementary Table 12**).

Considering the deep conservation of transcription factor recognition sites across metazoans, we next applied BOM to zebrafish data. Based on bulk ATAC-seq data across 11 tissue types, BOM classified 135,763 distal and non-exonic CREs with high performance (mean F1: 0.96, auPR: 0.99, auROC: 0.99; **Fig. 2c** and **Supplementary Table 13**).

Finally, in *Drosophila*, BOM discriminated between enhancers with either a development or housekeeping preference based on their activity as measured by STARR-seq under a housekeeping or development promoter. Development and housekeeping enhancers were previously defined in Drosophila S2 cells using STARR-seq ^61^ (**Supplementary Table 1**). Using a training set of enhancers ^61^ representing 15.6% of the data used for DeepSTARR ^53^ – a deep learning model trained on all genome-wide measurements, BOM achieved Pearson correlations of 0.78 and 0.42 for housekeeping and developmental promoters, respectively, compared to DeepSTARR’s 0.74 and 0.68 (**Fig. 2d**).

In summary, BOM demonstrated high overall performance, effectively learning the sets of context-specific motifs. This versatility underscores the potential of BOM for understanding the regulatory code across different biological systems.

### BOM uncovers the key motifs driving model prediction

In addition to identifying regulatory regions, we want to understand potential mechanistic driver genes to understand how cells select the repertoire of CREs necessary for maintaining their distinct identities and functions. We use SHAP (Shapley additive explanations), a model explanation approach rooted in cooperative game theory. SHAP is able to handle non-linear and interaction effects between motifs to interpret complex models. Using a tree-based SHAP method ^62^, the approach aggregates results across all CREs and calculates the marginal contribution of each motif to the classifier’s prediction. This contribution, an importance score, reflects how much a motif influences the prediction and can be used to provide local and global interpretability, where motif importance is defined by the average reduction in training loss gained when a motif is used to bifurcate a tree. The method provides the overall importance of each motif within the cell-state-specific model (global interpretation) and of individual motif contributions to specific CREs (local interpretation).

On average, ∼83.4% of total bases (500 bp) in distal elements were annotated by motifs with an importance score above 0 across all cell types in the mouse embryonic data ^47^. Per sequence, 32 motif instances have non-zero SHAP scores.

BOM explains the key motifs used in the cell state prediction for each regulatory element. To demonstrate this, we focus on three regulatory elements upstream of Nkx2-5, an early cardiac lineage regulator in embryonic mouse E8.25 (**Fig. 3a-c**). As expected, the sequences were predicted as cardiomyocyte-specific, consistent with Nkx2-5’s pivotal role in cardiac differentiation. Their associated motifs were linked to TFs with recognized roles in heart development, such as Mef2c and Srf. Interestingly, the motifs most important to these predictions differed between elements, highlighting potential functional heterogeneity between these adjacent sequences.

**Fig 3.**
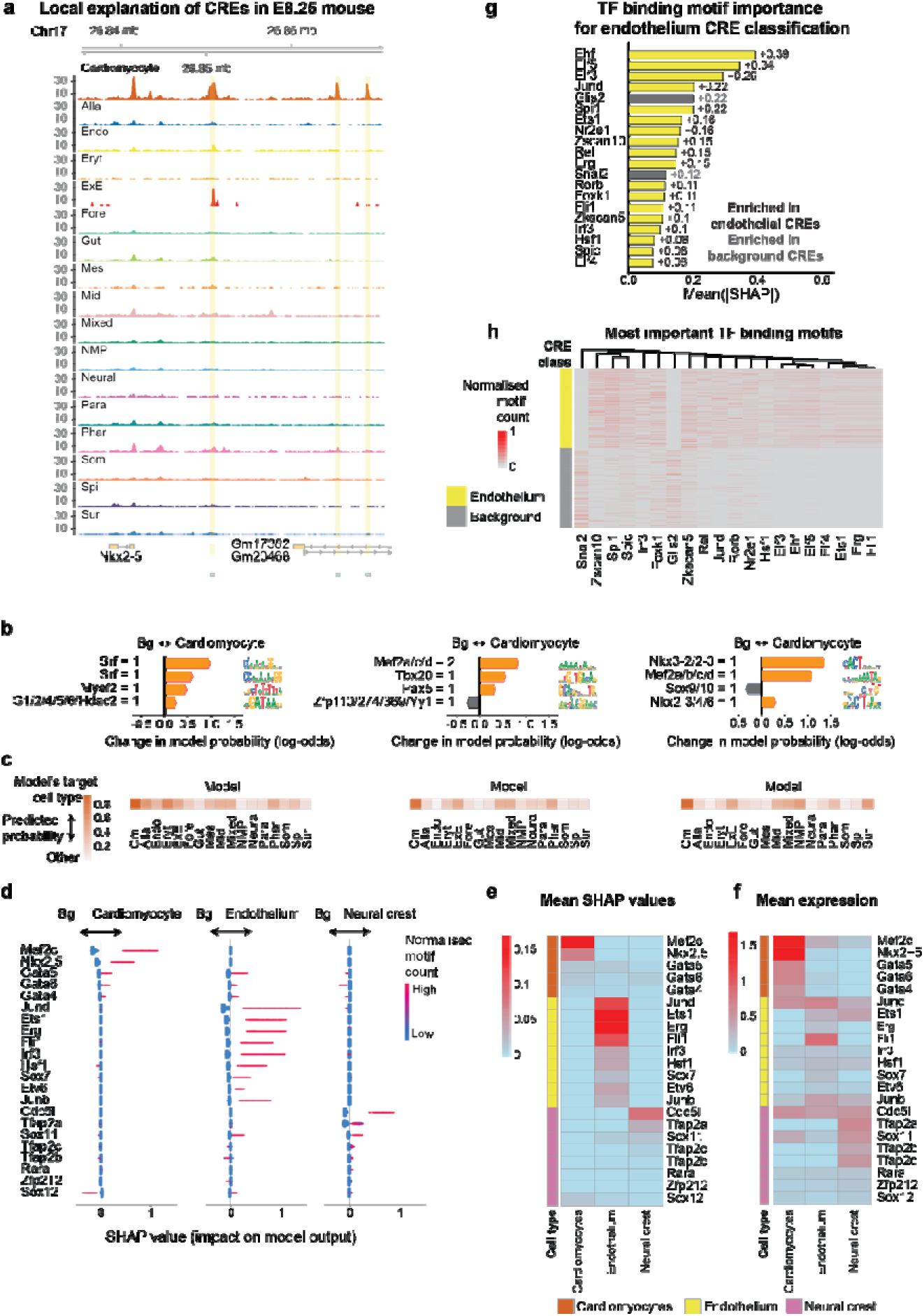
BOM provides local and global motif importance scores. **a.** Genome browser tracks showing the snATAC-seq signal around a region of mouse chromosome 17 near the gene Nkx2-5 for each mouse E8.25 cell type. The location of three cardiomyocyte-specific CRE are shown at the bottom. **b.** SHAP local explanation of three cardiomyocyte-specific CREs shown in **a.** The top four most important motifs for classifying those CREs are shown. The red and blue arrows indicate the sign (and the direction) of SHAP values. A representative name was given to every motif, and the motif count is indicated. **c.** Heatmaps representing the predicted probability of the CREs shown in **a** and **b** by each of the binary models trained to predict CREs specific to a cell type in mouse E8.25 **d**. SHAP values of top TF binding motifs in distinguishing cardiomyocyte-, endothelium-, and neural crest-associated CREs in mouse E8.25. The SHAP values were calculated based on background sets of CREs specific to other cell types. Each dot represents the SHAP value of a CRE for a given motif. The color code represents the normalized motif count: (counts - min(counts, na.rm = TRUE))/(max(counts, na.rm = TRUE) - min(counts, na.rm = TRUE))). A positive SHAP score indicates importance in the target CRE set, while a negative value indicates importance for the background set. **e**. Mean SHAP values for the TF binding motifs are shown in **d**. **f**. Mean expression of the TFs that bind to the motifs in **d** and **e**. Expression data are normalized counts from matched scRNA-seq experiment ^63^. **g**. Top 20 most important motifs in distinguishing mouse E8.25 endothelium CREs. Motifs are ranked by the mean of the absolute SHAP values. The bars are colored yellow or grey, representing positive and negative SHAP values, respectively. **h**. Normalized motif counts of the motifs shown in **g**. The motif counts normalized as **d**. The CREs shown are from the dataset used for endothelium CRE classification, where the background set is composed of CREs specific to the other 16 cell types in mouse E8.25 (n = 1760, 1768; background and endothelium CREs, respectively). Motifs were hierarchically clustered. Cis-BP 2.0 motifs for *M.musculus* in MEME Suite (v 5.4.0) were used in this figure for direct comparisons between motifs and TFs ^64^.

Where gene expression data is available, motifs can be associated with TF expression. We compared motif importance scores and gene expression to show that motif importance can reflect known biological drivers of cell identity and differentiation using matched scRNA-seq data for mouse gastrulation ^63^. We aggregated mean SHAP values across CREs to obtain overall importance per motif and matched the most important motifs to candidate TFs using Cis-BP motifs ^64^. Many TFs showed lineage-specific expression patterns reflective of the importance of their motif (**Fig. 3d-g**). For example, important TFs for cardiomyocytes included Mef2c, Nkx2-5, and Gata, while Jund, Ets1, Erg, Fli1, Irf3, Hsf1, Sox7, Etv6, and Junb are associated to the endothelium. We also observed clear distinctions in raw motif counts between the predicted cell type and the background set of CREs consistent with their motif importance scores (**Fig. 3h**).

### BOM motifs direct human cell-type-specific expression using synthetic regulatory elements

To validate BOM’s ability to identify key functional motifs, we asked whether we could create CREs with cell type specific enhancer activity based on engineering an enhancer sequence using motifs with the greatest explanatory power. To test this, we took the top five motifs ranked by global importance scores from HepG2 and Gm12878 models, which originate from liver cancer cells and blood lymphocytes, respectively, and constructed synthetic regulatory elements (SREs) by randomly inserting their respective motifs into the same muscle enhancer, serving as a sequence template ^43,65^. Five SREs were generated for each cell type, i.e., for each cell type, the same set of motifs were inserted at different locations and in random order. Using an enhancer activity reporter with a TATA minimal promoter, we tested enhancer activity of each sequence in both HepG2 and Gm12878 cells, normalizing for transfection efficiency and the basal activity of the template sequence.

We found cell-type specific SREs consistently drove the template sequence towards increased reporter expression in either HepG2 or Gm12878 cell lines depending on the inserted motifs (**Fig. 4**). Enhancer activity of the template sequence was low in both cell lines (**Supplementary File 1**). Interestingly, the level of transcription driven by SREs varied by 3 to 83 folds within cell-type specific SREs. Our results support that the type of regulatory motifs were the key determinants of cell-type-specific activity, while the order and spacing between the motifs tuned the strength of enhancer activity. We observed higher enhancer activity in the liver-derived HepG2 cell line over the Gm12878. As liver gene expression is known to be driven by a small number of TFs, this may explain the higher activity levels observed in HepG2 ^66^. In summary, our result demonstrated the ability of BOM to identify key motifs. It revealed that these sequences play a role in directing cell-type specific expression and can be used to generate cell-type specific synthetic elements.

**Fig. 4.**
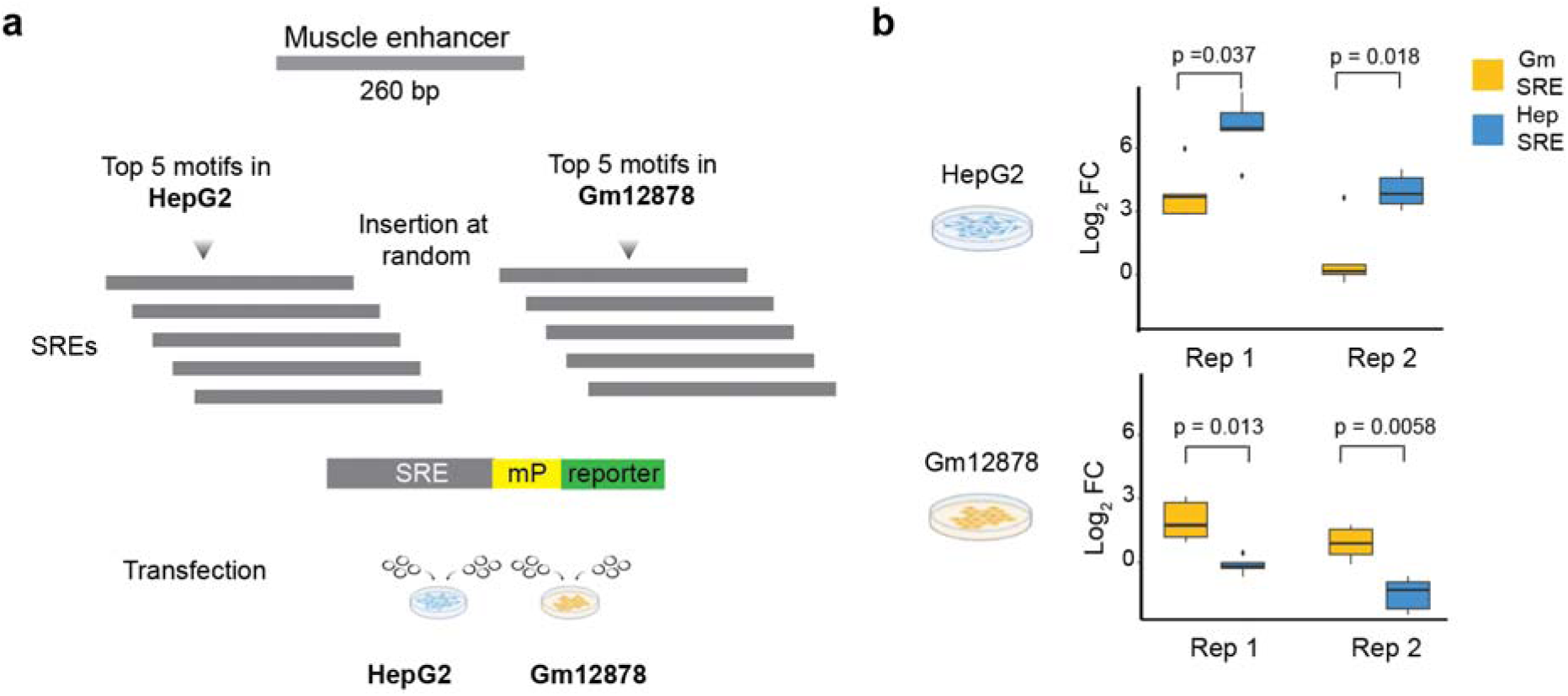
BOM identifies motifs that drive cell-type dependent activity. **a**. Top 5 ranked SHAP motifs for HepG2 and Gm12878 were implanted into a common template sequence (2 copies of the motif in each sequence) to construct synthetic enhancers (SREs)(HepG2 SRE n=5, Gm12878 SRE n=5). Each SRE was inserted into a reporter containing a minimal TATA promoter and transfected into both cell types **b**. Boxplot shows the log2 fold change of SREs accounting for transfection efficiency and the template enhancer activity. Two replicates of the assays were performed—P-values from two-tailed t-tests.

### BOM detects the evolutionary conservation of CREs

Enhancers exhibit a rapid rate of evolution. In a study of human liver candidate enhancers, only 1% displayed strong sequence conservation among placental mammals ^16^. This high evolutionary rate is advantageous in terms of mutational robustness, as the flexible nature of regulatory sequences allows them to maintain function despite significant sequence divergence ^15^. This flexibility, however, makes their identification using traditional sequence alignment methods challenging. To address this issue, we investigated whether BOM can capture generalizable sequence features between cell types in cross-species comparisons.

To compare species, we tested how well a human model trained to distinguish fetal human cardiomyocytes from cell-type associated CREs classified mouse embryonic cardiomyocytes CREs (**Fig. 5a**). Mouse cardiomyocytes were also compared with other cell types associated CREs. Results showed that the human model correctly predicted 64.5% of mouse CREs. Similarly, a model trained on human erythroblasts used to predict mouse erythroblasts CREs showed 62.9% accuracy (**Fig. 5a-c** and **Supplementary Table 14-15**). Of the correctly predicted CREs, 90.1% and 96.6% of cardiomyocyte and erythroblast putative enhancers could not be unaligned using LASTZ alignment between human and mouse genomes (UCSC liftOver, minMatch = 0.95%) (**Fig. 5d**). After relaxing alignment stringency to at least 60% sequence match across the length of the CREs, approximately a third (29.9% and 37.5%, respectively) of the CREs correctly classified by BOM remained unaligned (**Fig. 5d**). A similar trend was found for erythroblasts and results were consistent when a mouse model was trained and tested on human data (**Supplementary Fig. 8**). The results demonstrate BOM can capture evolutionary similarities based on collective motif composition beyond those detectable by conventional sequence alignment (**Supplementary Fig. 9)**. We expect similar sequence dependencies in other cell types across different species.

**Fig. 5.**
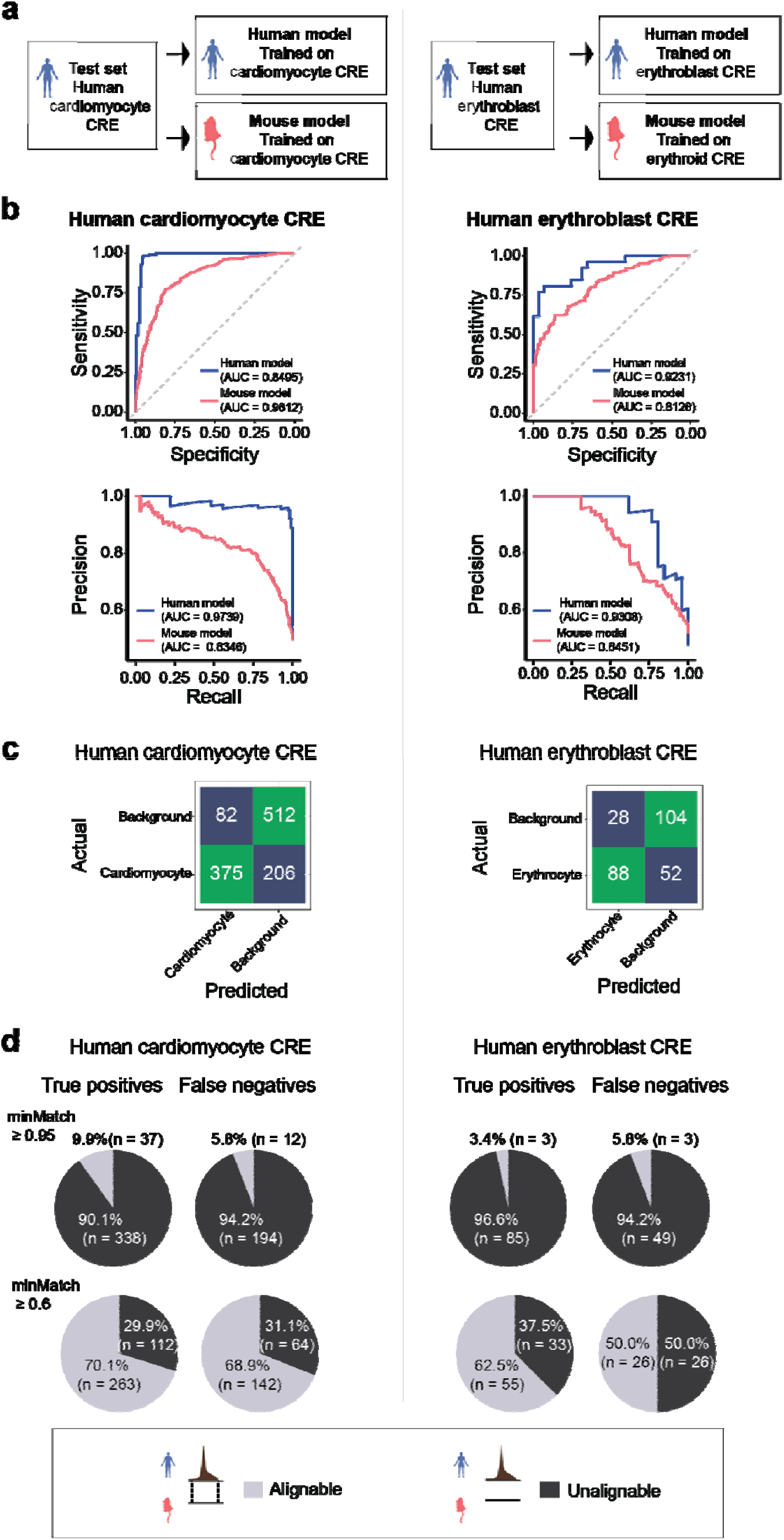
Cross-species prediction of cell type-specific CREs. **a**. Binary models were trained to distinguish cell type-specific human fetal CREs or mouse E8.25 CREs from a background set of CREs specific to other cell types. Both models were used to predict human CREs, an independent CRE set not used during training. **b**. ROC (top) and precision-recall (bottom) curves for the prediction of human fetal cardiomyocyte (left) and erythroblast (right) CREs using the models trained on human (blue) or mouse motif counts (pink). **c**. Confusion matrices for cardiomyocyte (left) and erythroblast (right) show the number of human CREs classified into their actual classes and the classes the models trained on mouse data predicted. **d**. Proportion of human cardiomyocyte (left) and erythroblast (right) CREs that can be aligned (gray) and cannot be aligned (black) to the mouse genome. The CREs were separated into the ones correctly predicted (true positives) and those incorrectly predicted as the background set (false negatives). The CREs were mapped to the mouse genome using liftOver with a threshold of minMatch >= 0.95 or 0.6 (top and bottom rows), where minMatch represents the minimum ratio of bases that should remap.

## Discussion

DNA sequences encode regulatory information in complex and flexible ways. Despite comprehensive genome sequences for various mammals and detailed maps of genomic features, the challenge of understanding distal elements that modulate spatiotemporal gene expression remains. We evaluated whether cell state can be quantitatively captured based on distal DNA sequences alone. This is analogous to asking whether cis-regulatory sequences contain sufficient sequence information to predict the cell type where it exhibits biochemical activity. To address this, we developed a ‘bag-of-motif’ approach for classifying and interpreting the cis-regulatory sequences of different cell states. Focusing on distal elements, known for their context dependency, we show that the count and combination of TF binding motifs are remarkably predictive of cell states in both embryonic and adult vertebrates. The model also performs well in modeling intermediary cell stages during embryonic development.

Hence, BOM is a predictive method to characterize the sequence basis of unknown distal regulatory regions. The ability to predict CREs using a held-out test set demonstrates that the motif combinations extracted from the training sequences were generalizable enough to predict previously unseen test sequences. This implies that distinct elements share characteristic sequence features and may function similarly. Our model’s performance, approaching 1.0, suggests that different cell types possess specific “codes” indicative of binding for various TFs. We validate our model prediction by creating human cell-type-specific synthetic enhancers, supporting the idea that these motifs can be sufficient in the cell types tested to drive gene expression in a cell type-aware manner. This demonstrates that motif composition is sufficient to predict cell type-specific regulatory activity in vertebrates. In contrast, the motif arrangement within an element tuned the strength of gene activity. It is well recognized that motif strength, orientation, and relative position can impose significant constraints, and the biochemical properties of DNA-binding protein occupancy and organization often require explicit motif organization ^67–69^. Our results are consistent with the ability of cis-regulatory sequences to recapitulate their endogenous regulatory states in a different trans-acting environment, even across species, as seen for DNA methylation ^70^, TF binding, and gene expression ^71^.

Beyond prediction, BOM explains each sequence based on the set of the most relevant TF motifs to the prediction. BOM helps to define the collectives of motifs to provide an integrated perspective of the cis-regulatory sequences that shape different cell states. We further demonstrate that BOM can classify cell type-specific enhancers that do not share genome-level alignment between humans and mice, illustrating the conservation of cell type-specific regulatory grammar across species, which has also been detected using deep learning models ^72–75^.

There are some caveats to our study. While we know that active cell-type specific regulatory regions are highly motif dependent, we have yet to determine how much of the remaining genome regions may share similar motif composition yet do not share regulatory activity. TFs do not bind all their motifs ^6^. Preferences for certain genome domains or nucleosome regions may restrict which motifs are bound and which regions are accessible ^76–78^. In regions of the genome less accessible to TFs, TF concentration may vary, which may lead to lower affinity of binding, which could particularly impact regulatory elements that are more dosage-sensitive ^79^. This could influence the types of CREs identified and impact BOM predictions. However, for most TFs, methods that measure open chromatin appear to be generally good predictors of *in vivo* binding ^5^. We also know that besides direct TF binding sites, nearby sequences can impact TF binding and context-dependent enhancer activity ^80–86^, motifs influencing DNA shape are also enriched in regions bound by TFs^84^. These sequence rules may be better captured using other strategies, such as k-mer enumeration ^36,87^ or deep neural networks (DNNs)(e.g. ^27,32,88^).

Finally, deep neural networks are powerful tools for genetic research, but we showed they can struggle to classify cell-type-specific regulatory elements at distal regions accurately. This is likely due to the limited amount of data in the genome to train the large number of cell states for DNNs and learn long range causal interactions ^89^. Based on our results, transfer learning of large DNNs may be a promising avenue to developing accurate representations of the regulatory contributions at distal elements. The multispecies datasets used here may offer a valuable resource for testing new models toward that goal.

## Methods

### Genome builds

The genome reference for mouse CREs is mm10, while the hg19 genome reference was used for human CREs. For fruit fly and zebrafish genome assemblies, dm3 and danRer10 were used, respectively.

### Defining candidate cis-regulatory elements

As described below, context-specific candidate regulatory elements were determined based on different experimental assays. Vertebrate regulatory elements must be over 1 kb from the transcription start site (TSS) and do not overlap exons. R packages ‘GenomicRanges’ and ‘GenomicFeatures’ (versions 1.42.0 and 1.42.3) were used for data processing ^90^. Numbers of CREs for each condition for the following datasets can be found in **Supplementary Table 1.**

#### Mouse E8.25 embryo

We used cell-type specific candidate cis-regulatory elements annotated based on scATAC-seq data from Pijuan-Sala et al. using Fisher’s exact tests ^47^ . Peaks were filtered out if annotated to multiple cell types or unannotated to any cell type. Peak numbers specific to notochord cells were low (n = 65 distal CREs) and were filtered out due to low peak count. CREs used as input were 500 bp by extending 250 bp from the middle of each peak. In addition to further capturing cellular heterogeneity in a cell type agnostic manner, we trained a classifier based on regulatory topics determined by cisTopic ^51^. CREs that were multi-label, where different topics existed for a given sequence, were removed. This resulted in a total of 76,604 CREs belonging to 93 topics.

#### Mouse E8.5 embryo

We obtained E8.5 marker ATAC-seq peaks from Argelaguet et al. ^56^. This dataset allowed us to test the performance of our models in an independent dataset that represented a similar developmental stage. To identify cell type-specific peaks, pairwise differential peak accessibility was performed using the negative binomial model with a quasi-likelihood test on five pseudo-bulk replicates in edgeR ^91^. Significant peaks were called with a 1% false discovery rate (FDR). Several criteria were applied to refine the selection of CRE candidates. We removed E8.5 CREs annotated as “Forebrain_Midbrain_Hindbrain” because, in the E8.25 dataset, the forebrain and mid-hindbrain are distinct groups. To ensure that test data was evaluated independently of training data, we excluded E8.5 regulatory regions that overlapped with elements used in training the E8.25 models from testing. Our final dataset comprised 12,079 CREs, each of 500 bp, centered on mid-peak.

#### Human cell lines

We collected 25-state ChromHMM models for six human cell lines (Gm12878, H1-hESC, Hela-S3, HepG2, Huvec, and K562) with hg19 genome reference from the UCSC Genome Browser ^60^. To define cell-line-specific CREs, we selected the regions annotated as strong enhancers in the ChromHMM models (states “Enh” and “EnhF”) and removed enhancers that overlapped strong enhancers in other cell lines. This resulted in a total of 167,148 candidate enhancers. All sequences were centered to 194 bp from the middle of each peak.

#### Human hematopoiesis

We used the single-cell healthy hematopoiesis chromatin accessibility atlas from Granja et al. ^92^. This dataset included scATAC-seq peaks annotated to 26 distinct blood and bone marrow cell types (n = 26,030). To define cell type-specific CREs, we adopted the strategy used by Pijuan-Sala et al.^47^. Initially, we assessed the accessibility of each peak across cells, counting the number of cells where the peak was accessible (count > 0) and the total number of cells for each cell type. We then conducted Fisher’s exact tests to determine the differential presence of peaks in each cell type compared to all other cell types combined (alternative = “greater”). Peaks not accessible in at least 5% of cells from the tested cell type were discarded. Fisher’s exact test p-values were adjusted with Bonferroni’s method for each scATAC-seq peak. Only the peaks with adjusted p-value < 0.01 were considered for further analysis. We then performed pairwise Fisher’s exact tests for each peak, comparing accessibility between pairs of cell types (alternative = ‘greater’). Again, peaks were tested only if present in at least 5% of cells within the tested cell type. Fisher’s exact tests p-values were adjusted with Bonferroni’s method for every scATAC-seq peak, and peaks with adjusted p-values > 1X10^-10^ in up to 13 pairwise cell-to-cell tests were retained for each cell type. Peaks annotated to multiple cell types were discarded, and the remaining peaks were defined as cell type-specific (n = 12,796). Human blood cell types “unknown 13” and “unknown 14” were excluded due to low element numbers (10 and 11, respectively). All CREs were 500 bp wide after extending ± 250 bp from mid-peak.

#### Fetal human

We used the human fetal chromatin accessibility atlas from Domcke et al. covering 15 organs, including the adrenal gland, cerebellum, cerebrum, eye, heart, intestine, kidney, liver, lung, muscle, pancreas, placenta, spleen, stomach, and thymus to define candidate CREs ^93^. To ensure cell-type specificity in the model, we excluded the cell mixture annotated as “Cardiomyocytes.Vascular_endothelial_cells”. Some fetal cell types are present in multiple organs. We aggregated these “multi-tissue” cells across organs for simplicity, although organ-specific differences may exist. These aggregated cell types include astrocytes, vascular endothelial cells, Schwann cells, stromal cells, myeloid cells, erythroblasts, lymphoid cells, smooth muscle cells, chromaffin cells, lymphatic endothelial cells, megakaryocytes, and ENS glia. Following this filtering and aggregation process, 71 cell types remained. To identify cell type-specific CREs, we implemented the same procedure described above based on Fisher’s exact test. The only difference was the final threshold, where peaks with adjusted p-values > 1X10^-10^ in as many as 20 pairwise tests were retained. A different threshold was selected due to the disparity in cell types between datasets. All sequences were centered to ± 250 bp from the peak center.

#### Adult zebrafish

We used zebrafish adult tissues ATAC-seq data from Yang et al. ^94^ to define CREs. The dataset includes peaks called in blood, brain, colon, heart, intestine, kidney, liver, muscle, skin, spleen, and testis. We filtered out overlapping ATAC-seq peaks for every tissue. Candidate regions (n=135,763) were 500 bp wide after extending ± 250 bp from mid-peak.

#### Fruit fly S2 enhancers

We obtained enhancers classified as housekeeping or developmental from STARR-seq enhancer activity assay in *Drosophila* S2 cells ^61^. STARR-seq was used to measure the ability of genomic regions to activate the ‘housekeeping’ core promoter RpS12 and the *Drosophila* synthetic core promoter (DSCP), a ‘developmental’ promoter ^53^. Enhancers were overlapped with the 200 bp regions tested by MPRA ^53^ and we calculated a mean value of “housekeeping” and “developmental” activity for each enhancer. The activity of the regions was measured as the log_2_FC over input (RNA versus DNA). After excluding enhancers used in training DeepSTARR, the final dataset we used comprised 6,287 enhancers of variable lengths ^61^.

### BOM Model

#### Motif database

We used the Gimme vertebrate TF binding motif database (v 5.0) to annotate CREs sequences (n = 1796 motifs) ^49^ (https://github.com/vanheeringen-lab/gimmemotifs). Motifs across multiple motif databases were clustered using agglomerative clustering, calculating motif similarity using Pearson correlation of motif scores >=0.5 to reduce redundancies. Although it comprises vertebrate motifs, it also performed well in *Drosophila,* presumably due to the deep conservation of TFs and their binding across metazoans ^13,95,96^. However, BOM is also compatible with enumerating with other PWM datasets (e.g., CIS-BP or JASPAR) or by gapped k-mer ^36^.

#### Motif scoring

*Motif instances were identified for each sequence* using FIMO from Meme Suite, with a threshold of 0.0001 (--thresh 0.0001) ^50^. FIMO scores the similarity between the input sequences and the motifs’ PWMs. FIMO incorporates a background model to determine the expected scores for random sequences lacking the motif. Our analysis applied a q-value cut-off of <= 0.5 to retain motif instances.

To investigate the impact of overlapping TF binding motifs in CRE sequences, we focused on mouse E8.25 CREs and generated a set of non-overlapping motifs for every cell type. To filter the motifs, we ordered them based on their start positions and systematically removed overlapping motifs with lower match scores until no motifs overlapped. When two overlapping motifs had equally highest scores, one was selected at random. Following removal, there was an average of 42.6 motifs per 1kb (q-value <= 0.5). Subsequently, we constructed motif count matrices and trained models using the abovementioned parameters.

#### Building classification data matrix

To create datasets for context-specific classification, we built a matrix of motif counts for each cell type where rows represent candidate CREs and columns represent TF binding motifs. We required a minimum of 100 unique peaks per context (cell type/condition/topic). For binary classification, every matrix contained a set of CREs of the target cell type (positive class) and a similar number of CREs from the rest of the cell types (negative class, balanced with stratified sampling to ensure an equal contribution from every non-target cell type). In cases where there are not enough CREs from a cell type to contribute to the background set, we lowered the contribution of every background cell type to that minimum number and adjusted the number of sequences from the target cell type.

### Training of classification models

#### Binary models

After building a matrix of motif counts to represent CREs, we randomly split the dataset into training, validation, and test sets (60%, 20%, and 20% sequences, respectively). To classify CREs, we used an Extreme Gradient Boosting (XGB) model due to its ability to handle high-dimensional datasets and nonlinear relationships ^46^. We used a set of hyper-parameters including nrounds = 10000, eta = 0.01, max_depth = 6, subsample = 0.5, colsample_bytree = 0.5, objective = “binary:logistic”, early_stopping_rounds = 100, eval_metric = “error” and maximize = F (xgboost R package version 1.6.0.1). We selected a small learning rate (‘eta’) to avoid overfitting, reflected in high performance on the test, unseen datasets. We used a maximum depth of the decision trees of 6 (‘max_depth’). A larger max_depth value produced negligible changes in the prediction performance metrics for mouse E8.25 cell type-specific CREs (comparison to max_depth = 8, 10, and 12) **(Supplementary Table 16)**. The fraction of the training instances to be sampled randomly in each boosting iteration = 0.5 (‘subsample’). The fraction of features sampled randomly for each tree = 0.5 (‘colsample_bytree’). We used the objective function ‘binary:logistic’ to train models for binary classification using logistic regression. We chose to minimize the training error rate. To prevent overfitting, if the error fails to decrease in 100 consecutive rounds (‘early_stopping_rounds’) or if the number of iterations reaches 10,000 (‘nrounds’), whichever is first, the training will terminate. We observed that in some cases, using a lower early_stopping_rounds value will result in the training stagnating in a local error minimum that was overcome by increasing the early_stopping_rounds value.

#### Multiclass models

For multiclass model training, we assigned the CREs to the same partitions as in the binary classification, ensuring consistency between the models. The same parameters were used in the multiclass models, except the objective function, which for multiclass classification was set to “multi:softprob”, and the eval_metric, which was set to “mlogloss” for multiclass classification error rate.

#### Training of regression models

We trained XGBoost regression models to predict housekeeping enhancer activity and developmental enhancer activity using enhancer activity data from STARR-seq ^61^. Similar to the classification models, we split the dataset into training, validation, and test sets, with proportions of 60%, 20%, and 20% respectively. The models were trained using the same parameters as the classification models, except for the objective function set to ‘reg:squarederror’, and the evaluation metric set to ‘rmse’. The objective function ‘reg:squarederror’ enables the models to optimize for accurate predictions in regression tasks, while the ‘rmse’ evaluation metric measures the root mean squared error to assess the prediction performance.

#### Prediction of CREs classes

Predictions were carried out with the function “predict” in all cases (R package stats version 4.0.0). The function returns an estimated predicted probability in the case of classification models and a predicted value of enhancer activity for the regression model. We applied a threshold of >0.5 for all classification models to define the predicted class.

### Model performance

#### Classification models

The performance of our classifiers was evaluated using multiple statistics. The key performance metrics are summarized as follows:

1. Precision: measures the model’s accuracy of positive predictions. TP / (TP + FP)
2. Recall: also known as sensitivity or true positive rate, measures the ability of the classifier to find all positive instances. TP / (TP + FN)
3. F1 score: the harmonic mean of precision and recall. 2 * ((precision * recall) / (precision + recall))
4. Matthews correlation coefficient (MCC)

In addition, we illustrate the performance of our models using Receiver Operating Characteristic (ROC) and precision-recall (PR) curves. auROC values were computed using the AUC function from the R package cvAUC (version 1.1.4). The auPR values were calculated using the pr_auc function from the R package yardstick (version 1.1.0).

#### Regression models

The models’ performance was assessed by calculating the Pearson correlation coefficient between the mean activity from S2 cells MPRAs data and the predicted activity values.

### Calculating motif importance

Shapley values ^97^ was used to quantify the contribution of each feature (i.e., motif) to the model’s prediction for each data instance (i.e., CREs). We calculate Shapley values using SHAP (Shapley additive explanations) ^98^. We train models with every combination of motifs as features. Once these models are trained with the same parameters and data, we predict a single observation using all of them. The difference in prediction when adding a motif represents its marginal contribution. The overall effect of a motif is then computed as the average marginal contribution across all models where the motif is included. These marginal contributions are aggregated using a weighted sum. The weights assigned to each model depend on the number of features included. The weight of a model with n features is calculated as one divided by the number of models with N features. This ensures that the sum of weights for all models with the same number of features equals 1.

Using our binary or multiclass classifiers and the training motif counts, we make an explainer object using the function shap.Explainer and calculate SHAP values with the function explainer (python SHAP v 0.41.0). A SHAP value is calculated for every motif and every CRE. We use rank motif importance using the sum of absolute SHAP values.

### Performance comparison

#### Binary models

BOM model performance was compared to two k-mer-based methods, LS-GKM and DNABERT ^36,52^. We trained LS-GKM and fine-tuned DNABERT on 17 datasets to predict mouse E8.25 cell type-specific CREs from background CREs. DNABERT models were fine-tuned on the same training and validation sets used in BOM models. LS-GKM models were trained on the union of the training and validation CREs used in BOM models, as it does not use a separate validation set. The input to LS-GKM is a file with the CRE sequences with a binary label. All peaks were extended to 500 bp regions from the peak summit in binary and multiclass model training. For DNABERT, CRE sequences were split into 6-mers using a custom script. LS-GKM was trained with default parameters, -t4 -l11 -k7 -d3. -t represents the kernel selection (wgkm), -l is the word length, -k is the number of informative positions, and -d is the maximum number of mismatches allowed. The 6-mer pre-trained DNABERT was fine-tuned to predict cell-type specific CREs for each cell type.

To fine-tune DNABERT models for each cell type, we first tested the recommended parameters: epochs=5, learning_rate=0.0002, logging_steps=100, warmup_percent=0.1, and weight_decay=0.01. During fine-tuning, we monitored the change in loss on validation sets and selected the model with the best loss for further evaluation. We also tested other learning rates, i.e., 0.0003 and 0.0001, and an epoch value of 20. We selected the model with the lowest loss for each cell type (**Supplementary Fig. 10**).

To fine-tune Enformer, an adaptive layer was added after the pointwise convolution layer while freezing all the weights in the previous layers. The adaptive layer consists of 2 fully connected linear layers with GELU as the activation function. The cropping layer of Enformer is modified to trim 704 positions on each side compared to 320 positions in the original Enformer paper, and our CRE is centered—the adaptive layer outputs two classes. Cross-entropy loss was used as the loss function. We performed hyperparameter searches for each cell type-specific model using Ray library (2.7.0) ^99^. Batch sizes between 4 and 8 were tested. The layer size was between 8, 16, and 32 to ensure any combination of batch size and layer size would not overfit CUDA memory. The Async Successive Halving ASHA ^100^ scheduler used the Bayesian Optimization HyperBand (BOHB) ^101^ search algorithm. Each cell type was tested with ten samples up to 10 epochs max. The checkpoint with the lowest loss was then used to test the model performance on the test set.

#### Multiclass models

For the multi-classification task, we trained models to predict 17 cell-type specific CREs and evaluated average performance metrics (auROC, auPR, and F1) across all cell types. For the multi-class prediction task, we constructed three popular deep learning architectures based on convolutional neural networks and hybrid architectures used by DeepMEL ^54^ & DanQ ^55^, DeepSTARR ^53^, and Basset ^28^; CNNx4+FCx2, CNN+LSTM, CNNx4+FCx2 CNNx3+FCx2, respectively (**Supplementary** Fig. 11). We re-implemented these three methods and trained the models on mouse development data to classify cell types. To adapt these methods to our task, we modified the output layer by setting the number of units (n) to 17, allowing the model to predict 17 cell-type specific CREs. We augmented the data by including their reverse complement sequences. Reverse complement was only performed for the CNN-based deep learning models, not BOM.

We used the same input and output configurations across all deep learning architectures. The input was represented using the one-hot encoding of 500 nucleotides (nt) sequences, while the output layer consisted of a fully connected layer with 17 units using a sigmoid activation function. We employed identical hyperparameters, including 100 epochs, a batch size of 128, early stopping of 10, and validation loss calculated using the ‘binary-crossentropy’ metric to train the models. We evaluated and selected the best model based on loss on the validation set.

To fine-tune Basset and DeepMEL, we reimplemented the Basset and the DeepMel models based on their respective datasets and published hyperparameters. We configured the training process with 60 epochs with an early stopping of five epochs. We froze pre-trained weights and fine-tuned the final output layer to tailor the model’s predictions to 17 distinct cell types. Ray library (2.7.0) ^99^ was used for a hyperparameter grid search. Learning rates for each model were methodically sampled from a uniform distribution, ranging from 1e^-5^ to 1e^-1,^ and batch sizes were evaluated at 64 and 128. The Bayesian Optimization HyperBand (BOHB)^101^ search algorithm and the Async Successive Halving ASHA scheduler ^100^ were used. Each model was tested with 40 samples up to 30 epochs max. The checkpoint with the lowest loss was then used to test the model performance on the test set (**Supplementary Table 5**). We performed a grid search on the original architecture trained on the mouse data for DeepSTARR but did not fine-tune it due to the degree of species divergence between insects and vertebrates.

### Cross-species CREs prediction

We trained binary models to predict human fetal cardiomyocytes and erythroblast CREs. We selected these models because of similar cell types in the mouse E8.25 dataset (cardiomyocytes and erythroid cells). The models were trained with similar parameters as in the mouse E8.25 models. Every model was used to predict elements in the same species and cross-species. Human and mouse CREs used in the cross-species analysis were aligned to the other species’ genome assembly using liftOver (v 435) with minMatch = 0.95 (default) and minMatch = 0.6. Chain files to carry out the alignment were obtained from UCSC for hg19 and mm10 assemblies.

### Gene expression data processing

*Mouse E8.25.* We used a single-cell RNA-seq mouse embryo time course atlas with gene expression spanning multiple developmental stages from E6.5 to E8.5 ^63^. Using the expression and metadata matrices, cells with less than 1,000 expressed genes, those with mitochondrial gene expression fraction higher than 2.37%, and those with mitochondrial read fraction outside their upper-end distribution were excluded, resulting in 116,312 cells ^102^. Expression was normalized using the function NormalizeData from Seurat (v 4.1.0)^102^ with default options (normalization.method = “LogNormalize”, scale. factor = 10000).

### Activity testing of synthetic regulatory elements

We selected the top five most explanatory motifs for HepG2 and Gm12878 from the ENCODE human cell line model we had constructed (**Supplementary File**). We used a muscle cell-specific enhancer as a sequence template (chr2:211,153,238-211,153,405), previously used for a similar purpose, and extended this to 260 bp (chr2:211,153,192-211,153,452) ^43,103^. We constructed 5 SREs for each cell line by inserting two copies of each motif in random order and location into the template to a final length of 260bp. We tested the 10 SRE and the template for enhancer activity in both cell lines using luciferase assay with a minimal promoter. Gm12878 cells (Corielle Institute of Medical Research) were cultured in RPMI media containing 15% FBS and 2 mM L-glutamine. HepG2 cells were gifted from N.Turner (VCCRI) and cultured in DMEM containing 10% FBS.

qPCR-based validation of the synthetic enhancer was performed as previously described ^104^. Briefly, the fragment to be tested was inserted upstream to the minimal TATA box promoter of the pGL4.10 (luc2) Vector (Promega #E6651). 5 million cells were then electroporated with the vector along with 2µg of a Renilla luciferase (RLuc) control plasmid (Promega #E2251). After electroporation, RNA was isolated after 6 hours, DNase treated, and reverse transcribed using primers shown below. qPCR was performed and analyzed using the delta-delta Ct method to quantify the expression levels accounting for transfection efficiency and activity of the muscle enhancer template in each cell type. Three technical replicates of qPCR were averaged. Two replications of the enhancer assays were performed using separate batches of cells.

FLuc F: CATTAAGAAGGGCCCAGC; FLuc R: GCTTCATAGCTTCTGCCAG

RLuc F: TAACTGGTCCGCAGTGGTG; RLuc R: TGGCACAACATGTCGCCA

## Code Availability

An R package and tutorial for BOM are available at https://github.com/ewonglab/BOM_package/. Code and data used for the manuscript are available at https://github.com/ewonglab/BOM_manuscript_data_scripts.

Processed datasets have been deposited to Zenodo (https://zenodo.org/records/10280300). Information on running times for BOM can be found in **Supplementary** Fig. 11.

## Acknowledgments

We thank R. Harvey (VCCRI), F. Zanini (UNSW), M. Francois (Centenary), and members of the Wong Lab for providing valuable critical feedback. We thank the Victor Chang Innovation Centre for support.

## Authors’ Contributions

P.C., X.Z., Z.L., D.H. performed computational experiments, data analyses, and code development. L.L. and Y.Y performed experimental assays and data analyses. P.C. and X.Z. contributed to manuscript preparation. E.W. conceived the study, designed experiments, analyzed data, and led manuscript preparation. All authors reviewed and approved the final version of the manuscript.

