## Supplemental text/figure for "A Bag-Of-Motif Model Captures Cell States at Distal Regulatory Sequences"

### Supplementary Figures

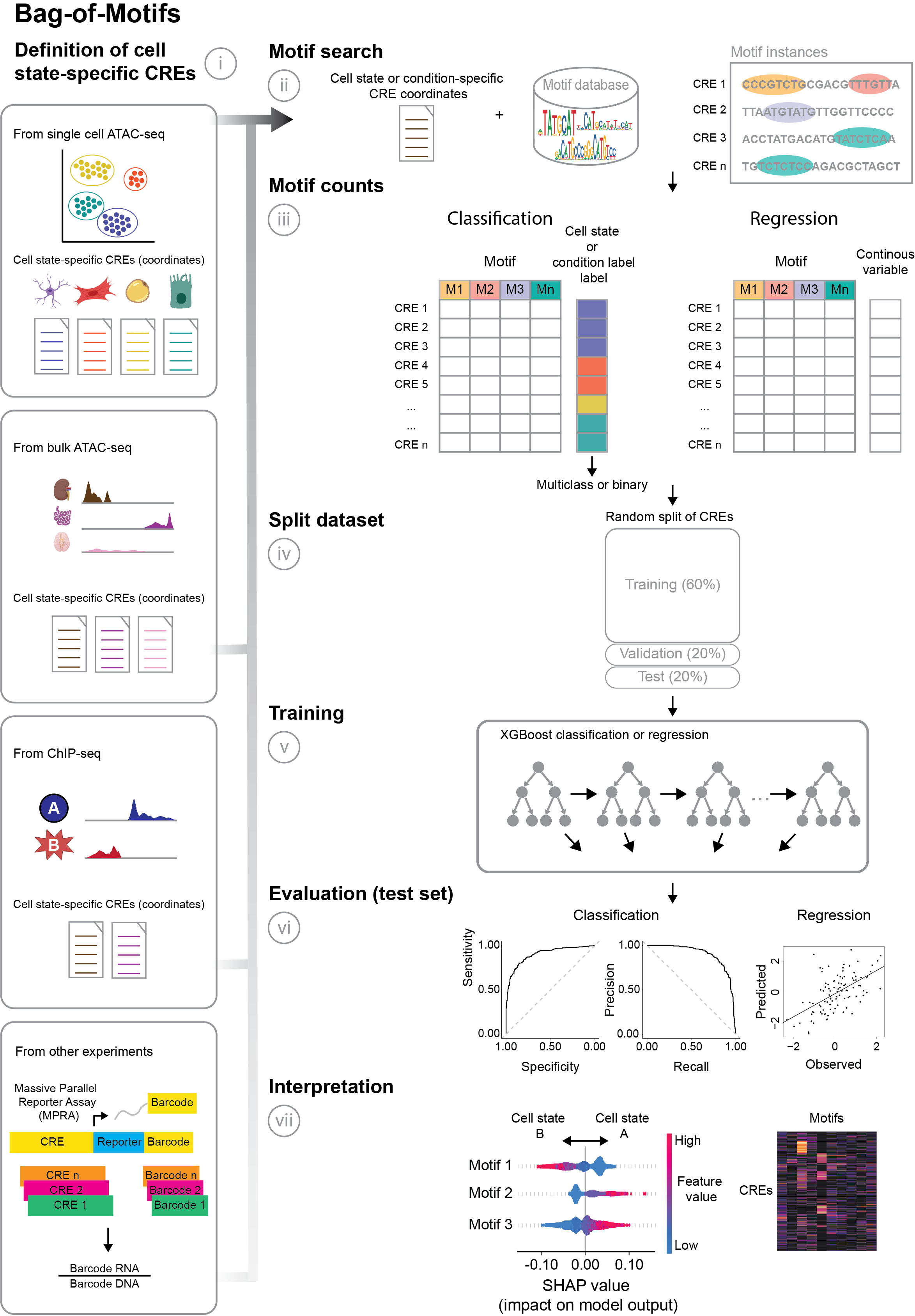

#### Supplementary Figure 1. A bag-of-motif (BOM) pipeline

The bag-of-motifs is a computational strategy for analyzing cis-regulatory elements (CREs) from different cell contexts. The pipeline can be applied to both classification and regression tasks and involves the following steps:

i. CRE Definition: CREs can be defined using different experimental data, such as single-cell or bulk ATAC-seq, or histone mark profiling (e.g., H3K27ac for active enhancers). These defined CREs are specific to particular cell states and can be labeled for binary or multiclass classification.
ii. TF Binding Motif Identification: FIMO ^1^ enumerates TF-binding motif instances within the defined CRE sequences based on motifs from PWM databases.
iii. Motif Frequency Matrix: A matrix is constructed with the frequency of motifs in each CRE. This matrix serves as input for subsequent model training.
iv. Dataset Splitting: The dataset is divided into training, validation, and test sets, with proportions of 60%, 20%, and 20%, respectively.
v. Model Training: An XGBoost ^2^ model is trained to predict target labels/values. Model performance is assessed during training on the validation set. Training terminates upon reaching a maximum number of iterations or when validation set performance ceases to improve after a specific number of iterations.
vi. Model Evaluation: The trained model's predictive performance is evaluated on the test dataset. Precision-recall curves and Receiver Operating Characteristic (ROC) curves are visualized for classification models. Higher values under the curves indicate better classification performance. For regression models, a correlation measure is calculated to assess prediction performance.
vii. SHAP Value Analysis: SHAP values are calculated to identify the TF binding motifs with the greatest influence on the predicted outcomes for each CRE.

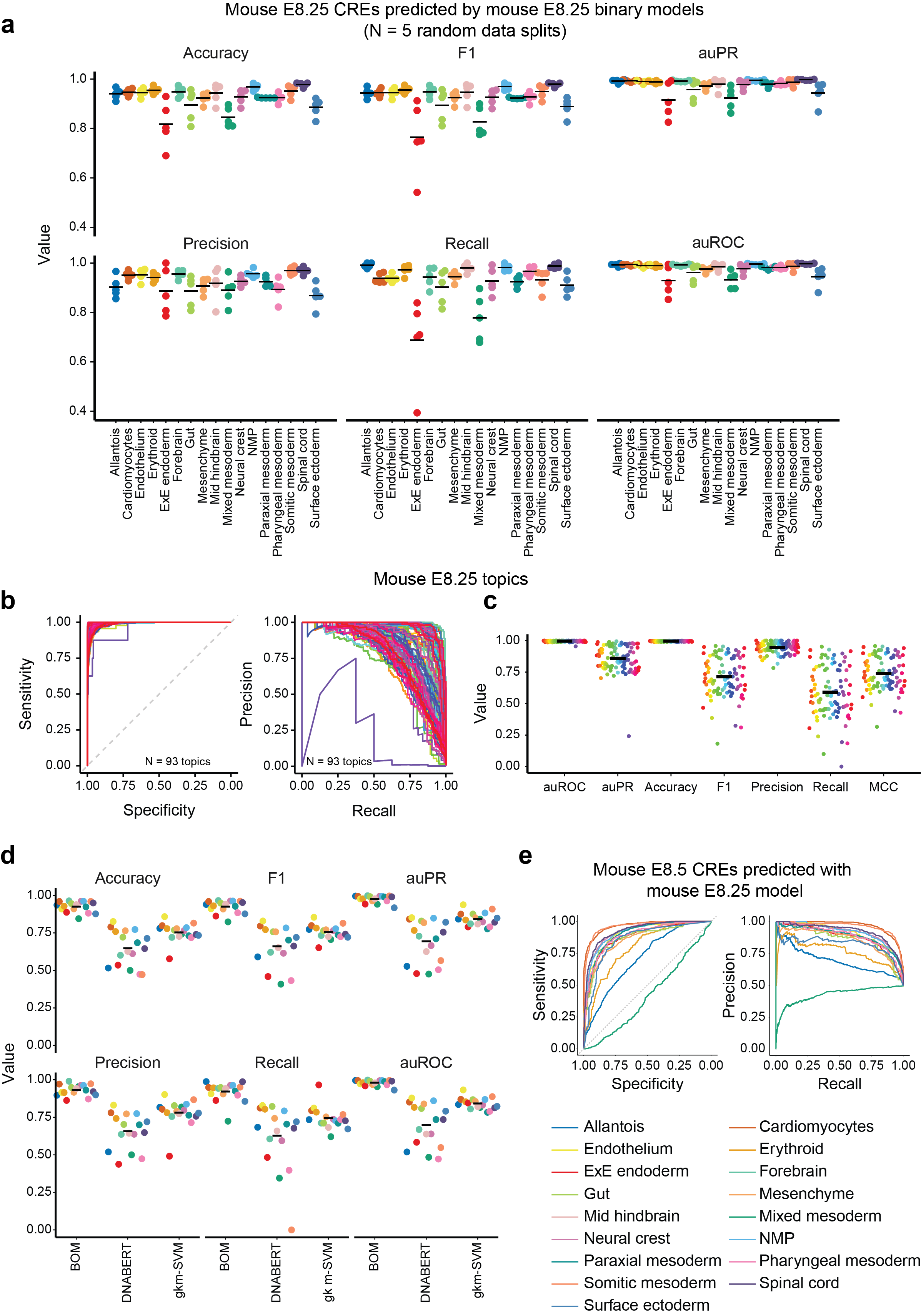

#### Supplementary Figure 2. BOM models for predicting mouse embryonic enhancers

**a**. Performance metrics for predicting cell type-specific enhancers in 17 mouse E8.25 cell types using binary BOM models ^3^. Accuracy, F1 score, area under the precision-recall curve (auPR), precision, recall, and are under the Receiver Operating Characteristic (ROC) curve (auROC) were calculated across 5 binary BOM models trained with different random data splits. Horizontal bars represent the mean value across the 5 test sets from the random data splits. The colors represent different cell types as in panels **d** and **e**. **b-c**. Prediction performance of multiclass BOM models trained to predict 93 mouse E8.25 topics across 15,321 test regions (**Methods**). **b**. ROC (left) and precision-recall (right) curves are shown. **c**. Performance metrics of models trained to predict mouse E8.25 topics. Area under the ROC curve (auROC), area under the precision-recall curve (auPR), accuracy, F1 score, precision and recall are shown. Horizontal bars indicate the mean value for every statistic. **d**. Performance metrics comparison for BOM, DNABERT and LS-GKM binary models trained to predict mouse E8.25 cell type-specific enhancers. Accuracy, F1 score, area under the precision-recall curve (auPR), precision, recall and area under the ROC (auROC) curve values are shown for the 17 cell types. Horizontal bars represent the mean value across the 5 test sets from the random data splits. **e**. ROC and precision-recall curves for predicting mouse E8.5 cell type-specific enhancers by models trained on mouse E8.25 enhancers. ROC curves (left) and precision-recall curves (right) are shown. The same color code is used for cell types in panels **a**, **d** and **e**.

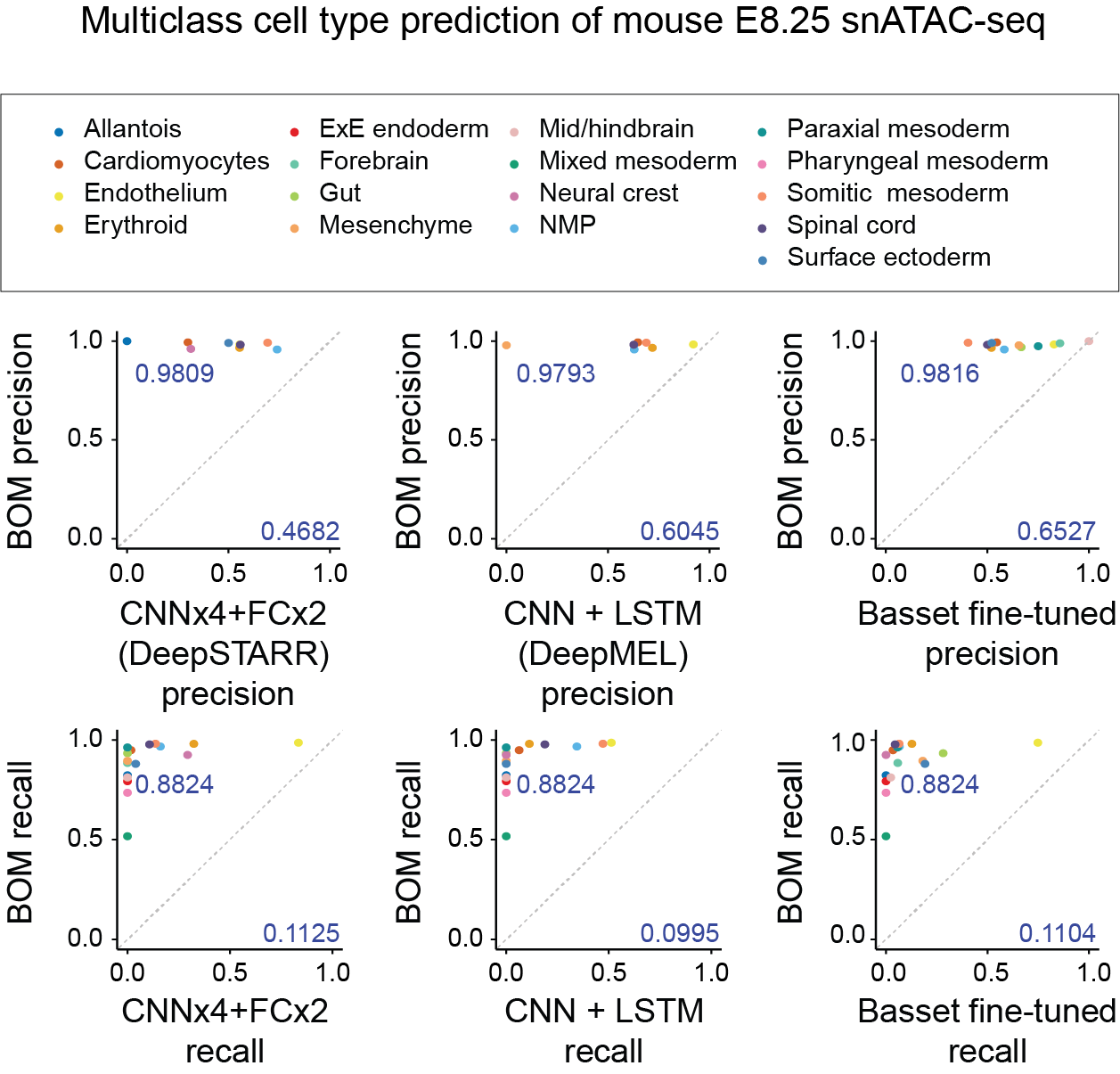

#### Supplementary Figure 3. Multiclass comparisons for predicting mouse embryonic enhancers

Performance metrics for predicting cell type-specific enhancers in 17 mouse E8.25 cell types comparing BOM to deep neural network architectures (DNNs) for multiclass predictions ^3^. These are common DNN architectures for sequence models in genomics. Hyperparameter searching and finetuning was performed as described in **Methods** and **Supplementary Table 6**.

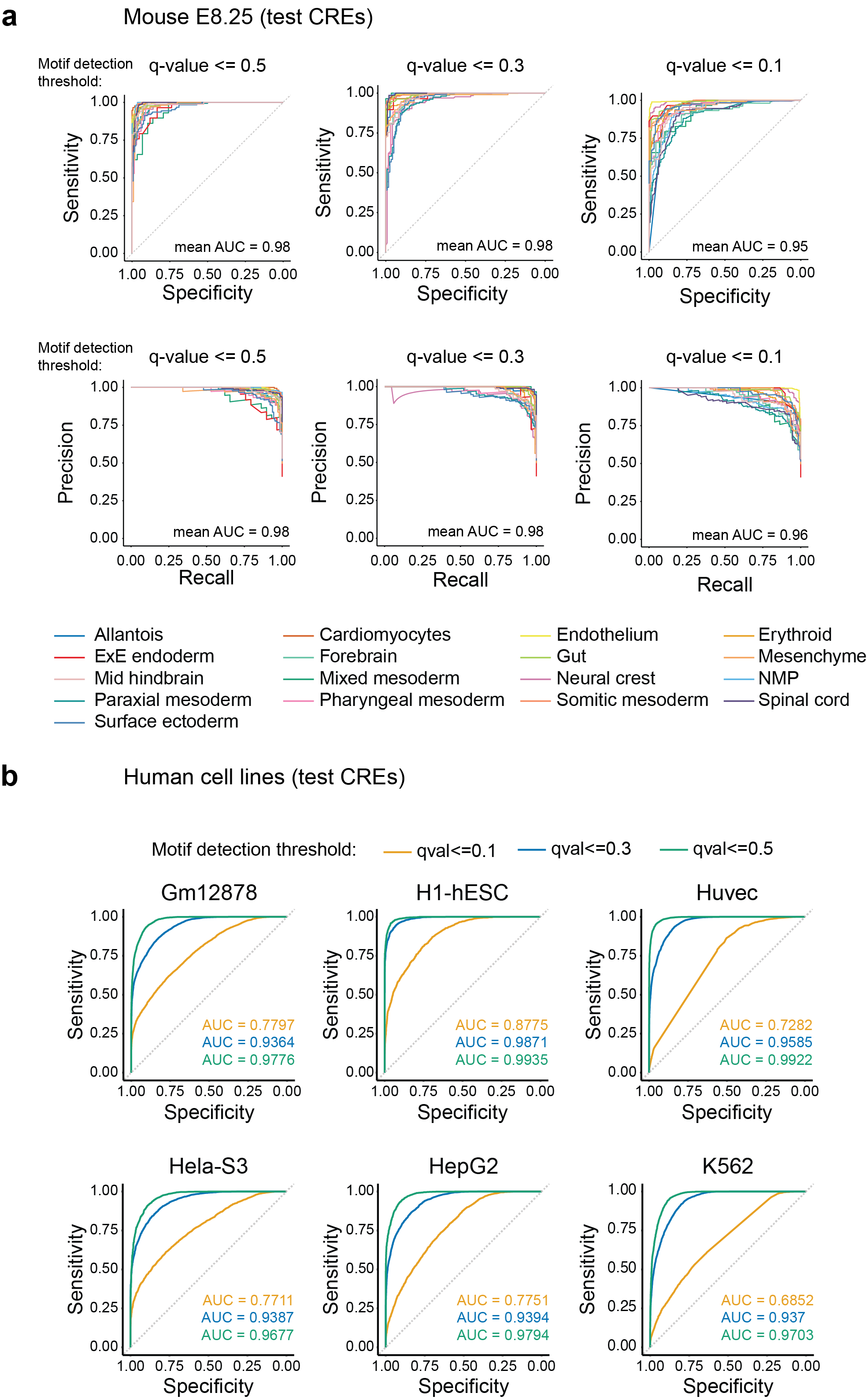

#### Supplementary Figure 4. Prediction performance for multiple motif detection thresholds

**a**. Performance of binary BOM models to distinguish mouse E8.25 cell type-specific enhancers across three motif detection thresholds (q-value <= 0.1, q-value <= 0.3, and q-value <= 0.5). Receiver Operating Characteristic (ROC) curves (top) and precision-recall (PR) curves (bottom) are shown. The mean area under the ROC curve (auROC) and area under the precision-recall curve (auPR) are shown for each q-value threshold. **b**. ROC curves are shown for six human cell lines: Gm12878, H1-hESC, Huvec, Hela-S3, HepG2 and K562. ROC curves for three motif detection thresholds are shown (q-value <= 0.1, q-value <= 0.3 and q-value <= 0.5). The corresponding auROC values are shown in each case.

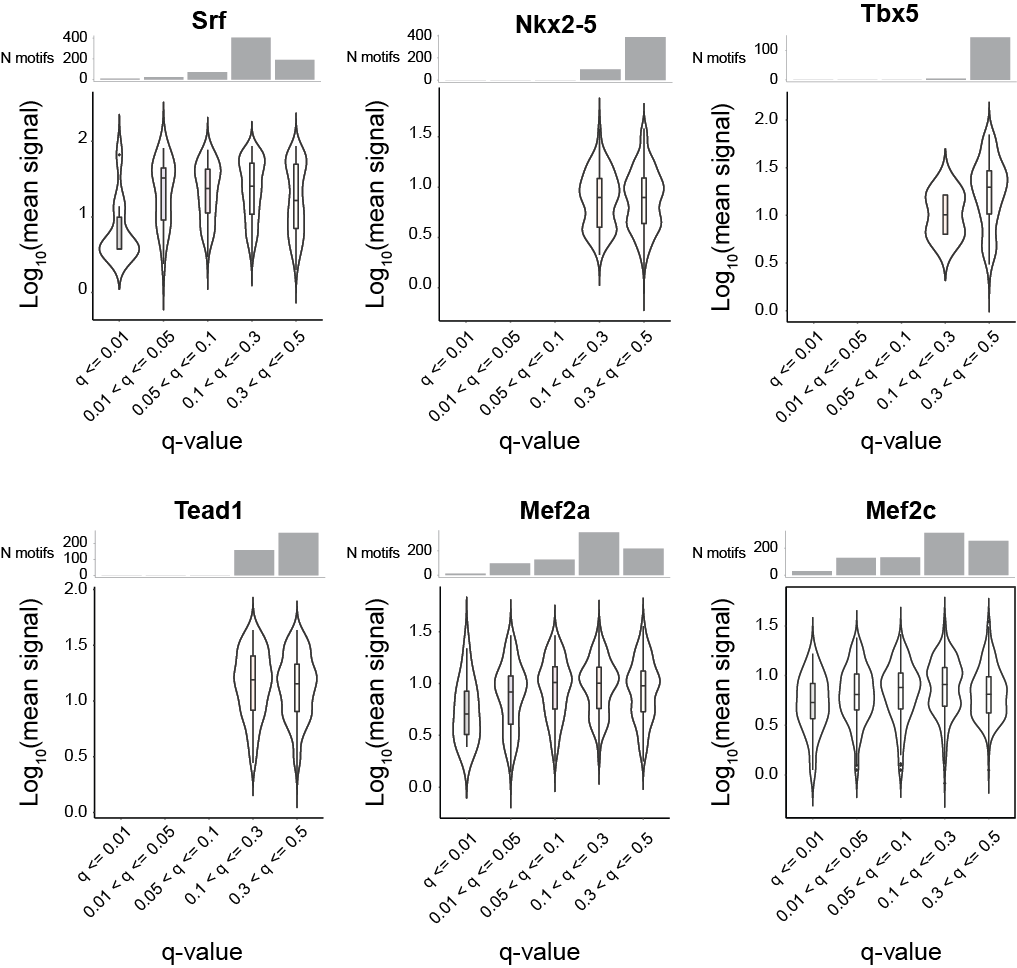

#### Supplementary Figure 5. TF binding sites are commonly missed using standard cut-offs for motif identification

TF binding signal using bioChIP-seq data compared to motifs of the corresponding TFs identified using FIMO ^1^ with the Gimme PWMs ^4^ at mouse embryonic CREs ^3^ for six cardiac developmental TFs (Mef2a, Mef2c, Nkx2-5, Srf, Tbx5, Tead) ^5^. The mean TF binding signal across each TF’s ChIP-seq summits are presented on the x-axis. Signals from the two biological replicates per TF were first averaged. TFs were matched using all corresponding motif PWMs.

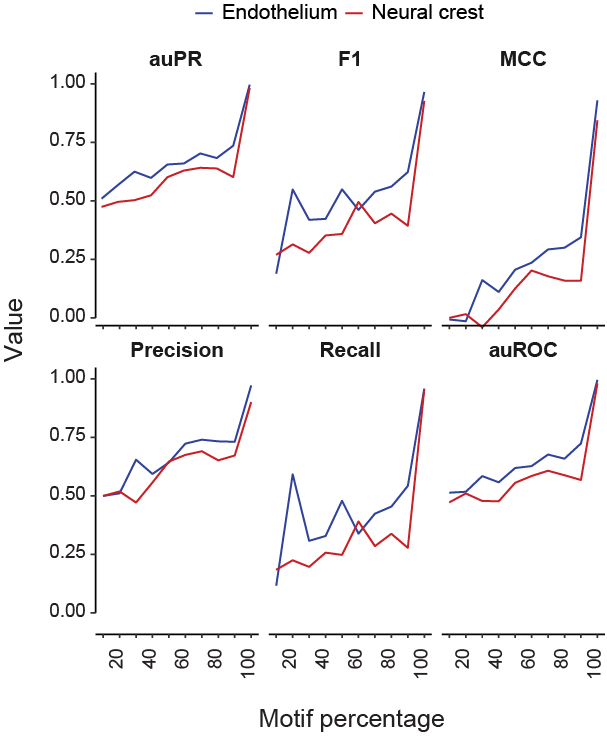

#### Supplementary Figure 6. BOM model performance dramatically decreases after random subsampling to reduce total motif counts

Performance of models trained for endothelium and neural crest distal CREs in the mouse embryonic test set.

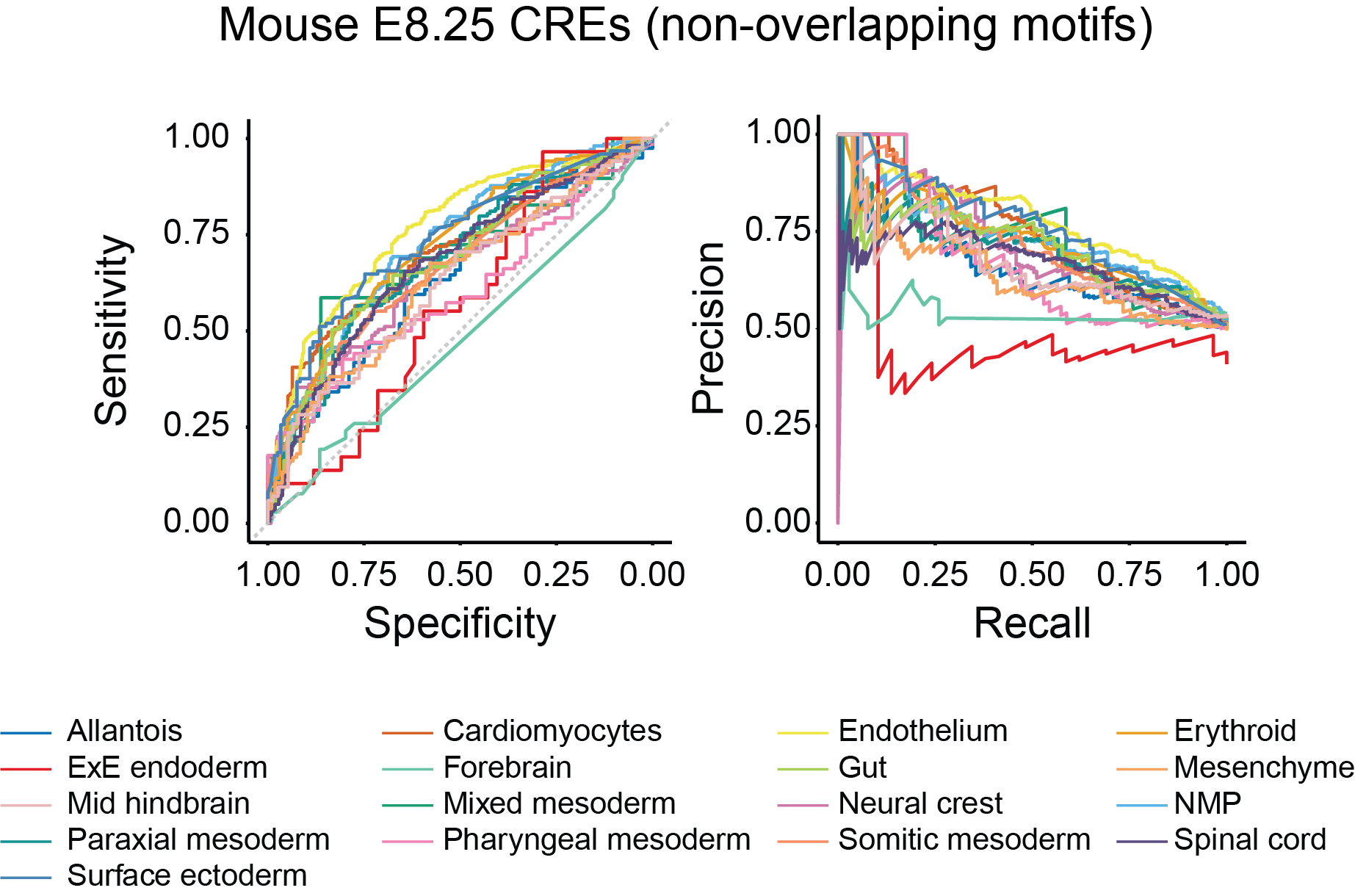

#### Supplementary Figure 7. BOM performance decreases if overlapping motifs are removed

Performance of models trained on non-overlapping motif counts. Non-overlapping motifs were defined as the ones with best motif scores; **Methods**). Receiver Operating Characteristic (ROC) curves (right) and precision-recall curves (left) are shown. A binary model was trained for each of the 17 mouse E8.25 cell types. The area under the ROC and precision-recall curves are shown in **Supplementary Table 10**.

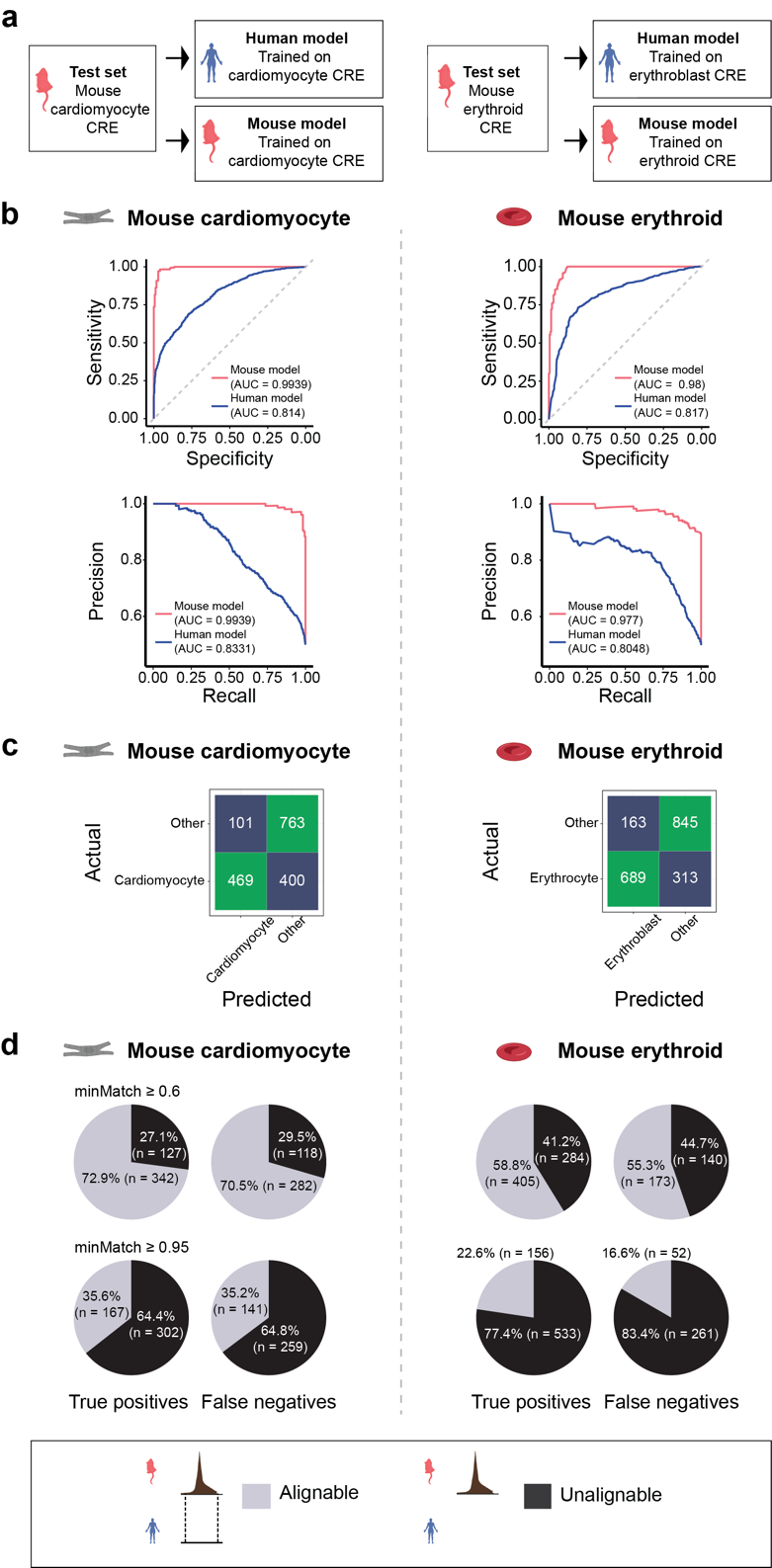

#### Supplementary Figure 8. Cross-species prediction of mouse E8.25 enhancers

**a**. Binary BOM models were trained to predict human fetal or mouse E8.25 cardiomyocyte or erythroid/erythroblast-specific enhancers. After training, the models trained on both species' data were utilized to predict mouse E8.25 enhancers. **b**. Top row: ROC curves for predicting cardiomyocyte (left) or erythroid (right) specific enhancers. The ROC curves obtained with BOM models trained on mouse data are depicted in pink, while the curves for predictions carried out with models trained on human data are shown in blue. The corresponding values of the area under the ROC curves are provided. Bottom row: PR curves for predicting mouse cardiomyocyte (left) or erythroid (right) specific enhancers using models trained on mouse or human data. **c**. Confusion matrices for predicting mouse cardiomyocyte (left) or erythroid (right) specific enhancers using models trained to predict human enhancers. **d**. Proportion of mouse cardiomyocyte (left) and erythroid (right) specific enhancers alignable (gray) or unalignable (black) to the human genome. Enhancers are categorized as true positives and false negatives. The top row of pie charts illustrates the proportion of enhancers alignable and not alignable using the threshold minMatch >= 0.6 in liftOver. The bottom row shows the proportions using the default threshold (minMatch >= 0.95).

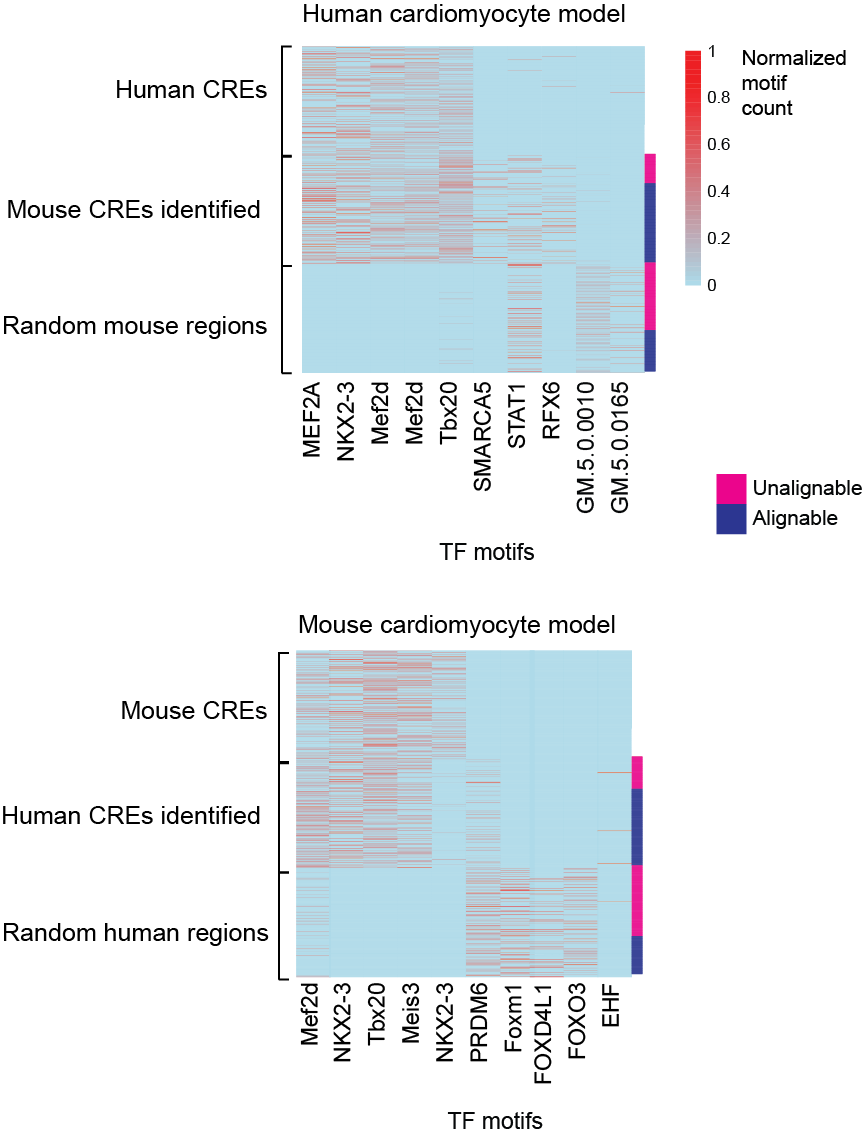

#### Supplementary Figure 9. The collection of motifs identifies CREs of similar cell types at similar developmental time points between human and mouse

Motif counts for human and mouse cell-type specific CREs for cardiomyocytes are compared to counts of sequences from the genome background. Human cardiomyocyte-specific CREs used to test the model trained using human cardiomyocytes are shown as the first rows in the top heatmap. Motif counts for accurately predicted mouse CREs are shown in the following rows. Random regions in the mouse genome of similar lengths are displayed in the bottom rows of the same heatmap. Motifs were chosen based on their ranked predictive values in predicting the positive cardiomyocyte class (top five motifs in columns 1-5) and the negative background (top five motifs in columns 6-10). The bottom heatmap shows the analogous information for a model trained on the mouse cardiomyocyte-specific CREs.

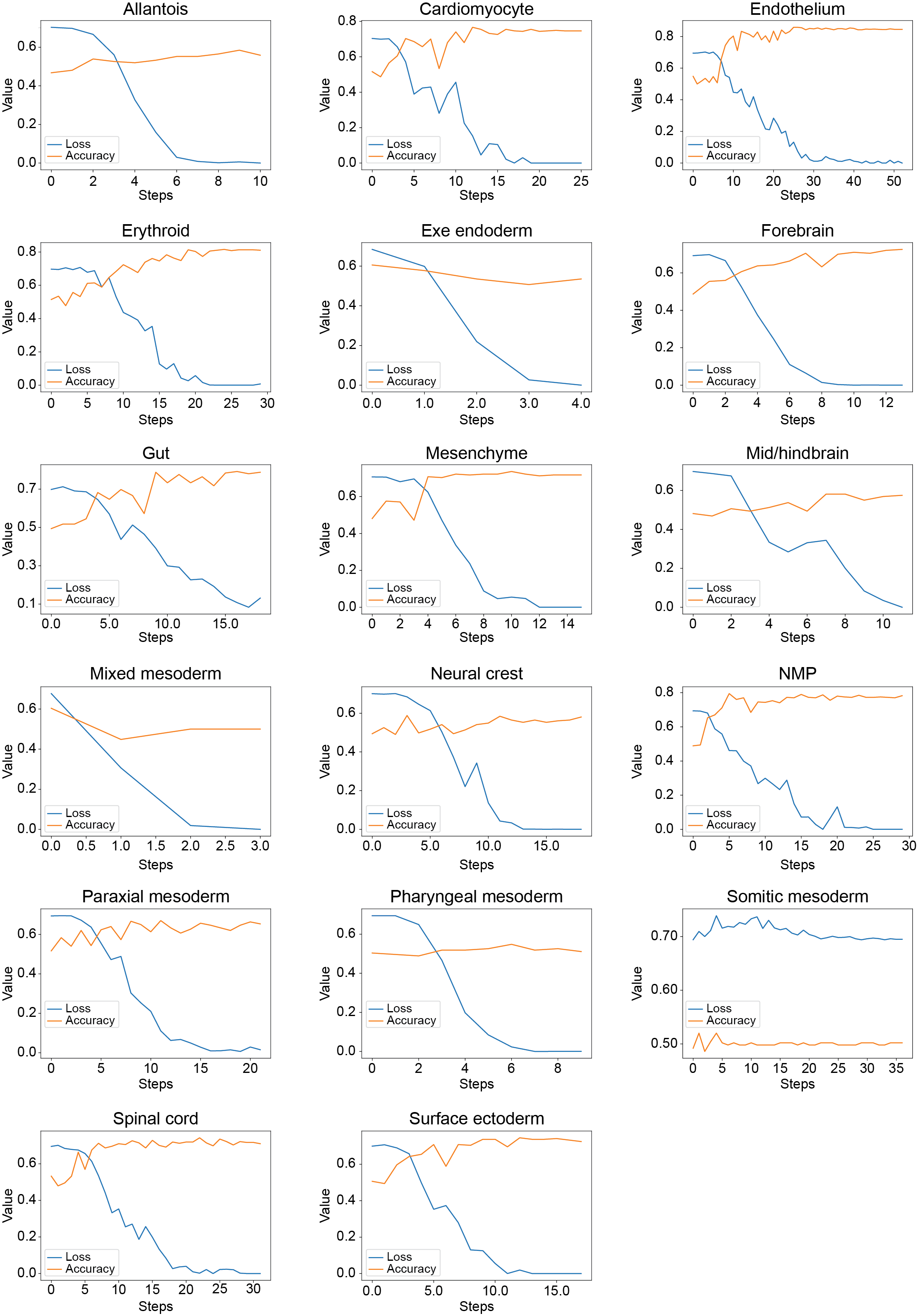

#### Supplementary Figure 10. Loss and accuracy across DNABERT training epochs

DNABERT models were trained to distinguish CREs specific to a cell type against a background set for each of the mouse E8.25 cell types. The loss and accuracy calculated on the validation set are shown across steps for each model.

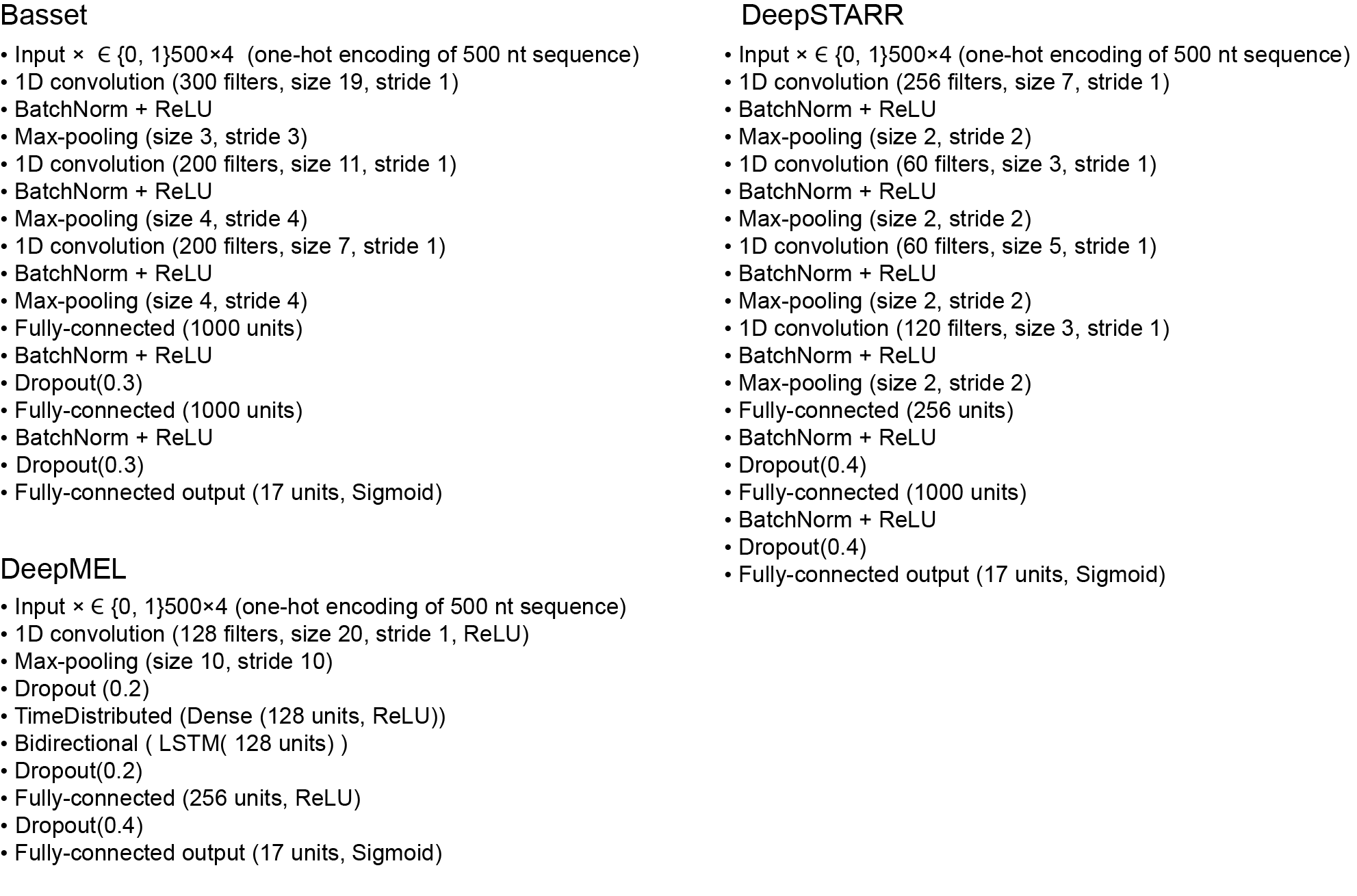

#### Supplementary Figure 11. Deep learning architectures

Three deep learning architectures are shown as used in Basset (CNNx3 + FCx2), DeepSTARR (CNNx4 + FCx2) and DeepMEL (CNN + LSTM).

**(a-c)** Processing time of motif annotation **(a)**, model training **(b)** and SHAP score calculation **(c)** for mouse E8.25 CREs. Motif annotation was carried out using a single CPU while model training and SHAP score calculation were executed in 4 CPUs. The X axis in **(a)** represents the number of cell-type-specific CREs per cell type (different colors represent different cell types as indicated in the box). In **(b)** and **(c)**, the x-axis the number of training instances (target and background) in each subset per cell type. Best fit line across the data points is shown in red. Shaded region represents the 95% confidence interval of the line of best fit. **(d-f)** Similar to **(a-c)** for human cell lines. The CPU time is shown in seconds for mouse CREs and in minutes for human CREs.

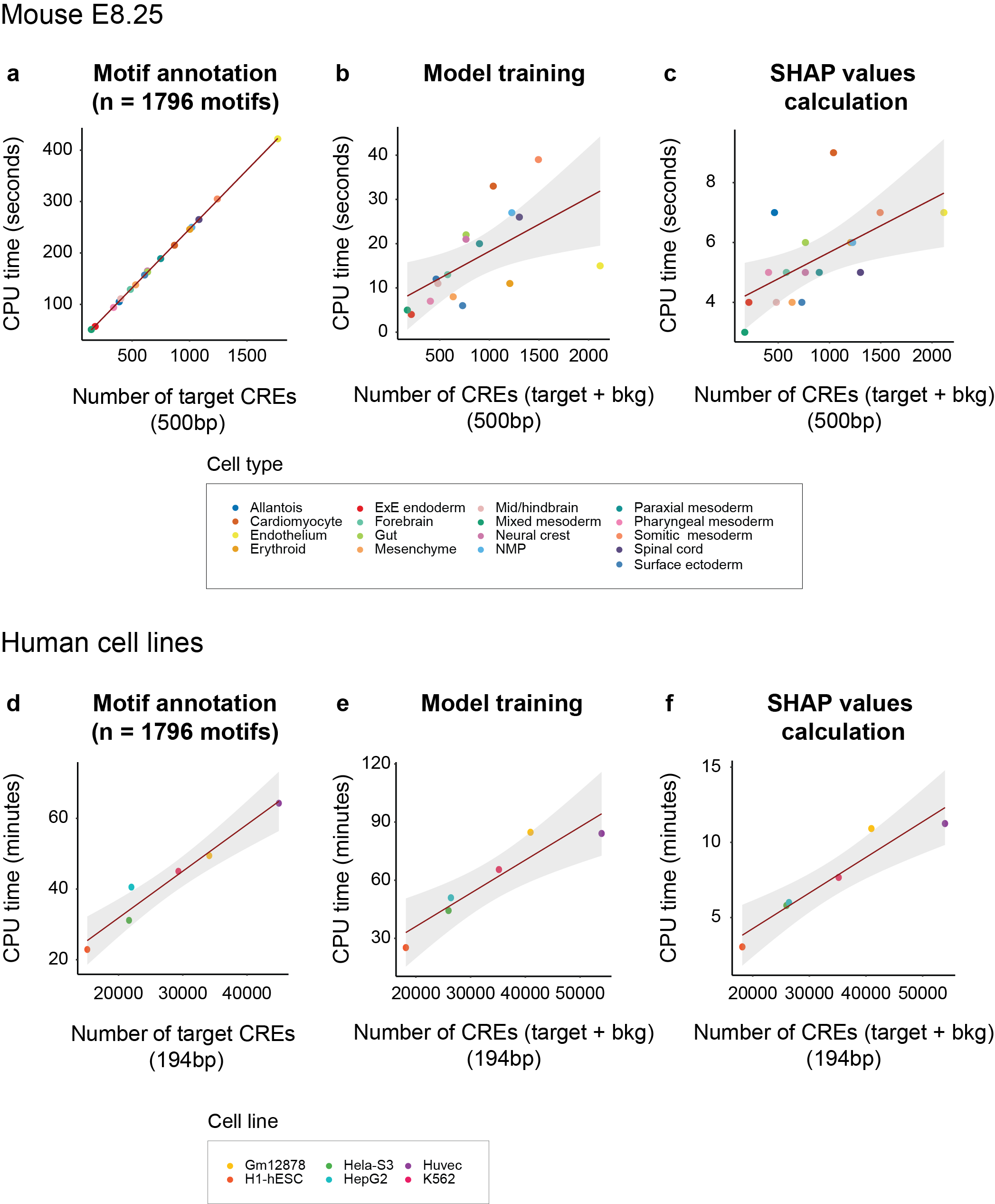

#### Supplementary Figure 12. BOM running times across datasets

**(a-c)** Processing time of motif annotation **(a)**, model training **(b)** and SHAP score calculation **(c)** for mouse E8.25 CREs ^3^. Motif annotation was carried out using a single CPU while model training and SHAP score calculation were executed in 4 CPUs. The x-axis in **(a)** represents the number of cell-type-specific CREs per cell type (different colors represent different cell types as indicated in the box). In **(b)** and **(c)**, the x-axis the number of training instances (target and background) in each subset per cell type. Best fit line across the data points is shown in red. Shaded region represents the 95% confidence interval of the line of best fit. **(d-f)** Similar to **(a-c)** for human cell lines ^6^. The CPU time is shown in seconds for mouse CREs and in minutes for human CREs.

#### Supplementary Table 1. Number of CREs used in BOM models

Number of cell types, cell lines, and condition-specific CREs used in BOM models. The numbers of training, validation, and test CREs are provided. The type of BOM model is indicated (binary, multiclass or regression). For binary models, the number of CREs represents the number of CREs for the target cell type of condition. For the multiclass model, the number of CREs represents the contribution from every “class” (e.g., cell type) to the dataset.

| Dataset  / Model type | Cell type (condition) | N CREs (training) | N CREs (validation) | N CREs (test) | total |
| --- | --- | --- | --- | --- | --- |
| Mouse (E8.25)  / Binary  /multiclass | Allantois | 235 | 74 | 79 | 388 |
|  | Erythroid | 606 | 192 | 204 | 1002 |
|  | ExE endoderm | 109 | 41 | 29 | 179 |
|  | Gut | 371 | 129 | 135 | 635 |
|  | mesenchyme | 314 | 112 | 105 | 531 |
|  | NMP | 608 | 200 | 212 | 1020 |
|  | Paraxial mesoderm | 445 | 145 | 159 | 749 |
|  | Pharyngeal mesoderm | 205 | 66 | 68 | 339 |
|  | Somitic mesoderm | 731 | 248 | 263 | 1242 |
|  | Spinal cord | 652 | 207 | 223 | 1082 |
|  | Surface ectoderm | 361 | 123 | 125 | 609 |
|  | Cardiomyocyte | 516 | 178 | 175 | 869 |
|  | Endothelium | 1059 | 348 | 361 | 1768 |
|  | Forebrain | 273 | 108 | 104 | 485 |
|  | Mid/hindbrain | 234 | 83 | 85 | 402 |
|  | Neural crest | 372 | 129 | 133 | 634 |
|  | Mixed mesoderm | 82 | 34 | 29 | 145 |
| Fruit fly S2 cells / Regression | Housekeeping and developmental enhancer activity | 3772 | 1257 | 1258 | 6287 |
| Zebrafish (adult) / Binary | Blood | 7896 | 2582 | 2627 | 13105 |
|  | Intestine | 5931 | 1964 | 2018 | 9913 |
|  | Skin | 18763 | 6258 | 6216 | 31237 |
|  | Brain | 12514 | 4182 | 4180 | 20876 |
|  | Kidney | 7311 | 2393 | 2439 | 12143 |
|  | Spleen | 3732 | 1278 | 1276 | 6286 |
|  | Colon | 2997 | 997 | 1008 | 5002 |
|  | liver | 5084 | 1652 | 1690 | 8426 |
|  | testis | 12505 | 4170 | 4173 | 20848 |
|  | heart | 8301 | 2736 | 2747 | 13784 |
|  | muscle | 4366 | 1429 | 1453 | 7248 |
| Human (fetal)  / Binary | Cardiomyocyte | 344 | 117 | 120 | 581 |
|  | Erythroblasts | 82 | 32 | 26 | 140 |
| Human  (cell lines)  / Binary | Gm12878 | 20483 | 6862 | 6776 | 34121 |
|  | H1-hESC | 9158 | 2974 | 3001 | 15133 |
|  | Hela-S3 | 12998 | 4349 | 4284 | 21631 |
|  | HepG2 | 13196 | 4428 | 4377 | 22001 |
|  | Huvec | 26926 | 9122 | 8903 | 44951 |
|  | K562 | 17629 | 5883 | 5799 | 29311 |
| Human (hematopoiesis) / Binary | B | 484 | 159 | 168 | 811 |
|  | CD14 monocyte 1 | 508 | 170 | 173 | 851 |
|  | CD14 monocyte 2 | 296 | 110 | 101 | 507 |
|  | CD4 M | 85 | 33 | 25 | 143 |
|  | CD4 N2 | 70 | 23 | 26 | 119 |
|  | CD8.CM | 228 | 77 | 82 | 387 |
|  | CD8.EM | 182 | 59 | 64 | 305 |
|  | cDC | 334 | 109 | 116 | 559 |
|  | CLP 1 | 392 | 134 | 143 | 669 |
|  | CLP 2 | 508 | 170 | 173 | 851 |
|  | CMP/LMPP | 295 | 107 | 100 | 502 |
|  | Early basophil | 508 | 170 | 173 | 851 |
|  | Early erythroid | 508 | 170 | 173 | 851 |
|  | GMP | 508 | 170 | 173 | 851 |
|  | GMP/Neutrophil | 508 | 170 | 173 | 851 |
|  | HSC | 135 | 47 | 44 | 226 |
|  | Late erythroid | 64 | 22 | 17 | 103 |
|  | NK | 420 | 141 | 153 | 714 |
|  | pDC | 508 | 170 | 173 | 851 |
|  | Plasma | 243 | 87 | 90 | 420 |
|  | Pre B | 508 | 170 | 173 | 851 |
|  | Unknown 26 | 310 | 110 | 103 | 523 |
| Mouse topics (E8.25)  / Multiclass | Topic2 | 741 | 243 | 237 | 1221 |
|  | Topic3 | 734 | 240 | 245 | 1219 |
|  | Topic4 | 56 | 23 | 20 | 99 |
|  | Topic6 | 223 | 63 | 65 | 351 |
|  | Topic7 | 377 | 120 | 137 | 634 |
|  | Topic8 | 255 | 110 | 113 | 478 |
|  | Topic9 | 186 | 71 | 87 | 344 |
|  | Topic10 | 181 | 78 | 69 | 328 |
|  | Topic11 | 684 | 192 | 212 | 1088 |
|  | Topic12 | 614 | 208 | 221 | 1043 |
|  | Topic14 | 371 | 137 | 115 | 623 |
|  | Topic16 | 422 | 151 | 147 | 720 |
|  | Topic17 | 269 | 105 | 98 | 472 |
|  | Topic18 | 567 | 162 | 188 | 917 |
|  | Topic19 | 548 | 177 | 192 | 917 |
|  | Topic20 | 364 | 152 | 139 | 655 |
|  | Topic21 | 513 | 165 | 141 | 819 |
|  | Topic22 | 235 | 83 | 63 | 381 |
|  | Topic23 | 812 | 240 | 266 | 1318 |
|  | Topic24 | 712 | 235 | 246 | 1193 |
|  | Topic25 | 575 | 173 | 158 | 906 |
|  | Topic26 | 503 | 161 | 201 | 865 |
|  | Topic27 | 307 | 96 | 104 | 507 |
|  | Topic28 | 782 | 248 | 282 | 1312 |
|  | Topic29 | 153 | 53 | 46 | 252 |
|  | Topic30 | 621 | 191 | 200 | 1012 |
|  | Topic31 | 313 | 128 | 120 | 561 |
|  | Topic32 | 760 | 271 | 239 | 1270 |
|  | Topic33 | 392 | 110 | 112 | 614 |
|  | Topic34 | 673 | 225 | 251 | 1149 |
|  | Topic35 | 491 | 176 | 163 | 830 |
|  | Topic36 | 754 | 251 | 247 | 1252 |
|  | Topic37 | 590 | 164 | 162 | 916 |
|  | Topic38 | 337 | 110 | 100 | 547 |
|  | Topic39 | 760 | 267 | 271 | 1298 |
|  | Topic40 | 186 | 78 | 57 | 321 |
|  | Topic41 | 445 | 132 | 147 | 724 |
|  | Topic42 | 383 | 105 | 136 | 624 |
|  | Topic43 | 491 | 158 | 172 | 821 |
|  | Topic44 | 577 | 203 | 197 | 977 |
|  | Topic45 | 756 | 237 | 251 | 1244 |
|  | Topic46 | 269 | 89 | 81 | 439 |
|  | Topic47 | 259 | 84 | 92 | 435 |
|  | Topic48 | 416 | 141 | 146 | 703 |
|  | Topic49 | 148 | 50 | 46 | 244 |
|  | Topic50 | 616 | 203 | 206 | 1025 |
|  | Topic51 | 439 | 145 | 153 | 737 |
|  | Topic52 | 569 | 179 | 214 | 962 |
|  | Topic53 | 315 | 121 | 107 | 543 |
|  | Topic54 | 740 | 255 | 255 | 1250 |
|  | Topic55 | 648 | 212 | 198 | 1058 |
|  | Topic56 | 773 | 253 | 236 | 1262 |
|  | Topic57 | 358 | 143 | 129 | 630 |
|  | Topic58 | 499 | 151 | 175 | 825 |
|  | Topic59 | 407 | 158 | 140 | 705 |
|  | Topic60 | 727 | 260 | 234 | 1221 |
|  | Topic61 | 439 | 144 | 140 | 723 |
|  | Topic62 | 429 | 149 | 127 | 705 |
|  | Topic63 | 474 | 178 | 158 | 810 |
|  | Topic64 | 687 | 207 | 237 | 1131 |
|  | Topic65 | 658 | 223 | 216 | 1097 |
|  | Topic66 | 779 | 291 | 286 | 1356 |
|  | Topic68 | 524 | 155 | 191 | 870 |
|  | Topic69 | 683 | 261 | 228 | 1172 |
|  | Topic70 | 783 | 252 | 231 | 1266 |
|  | Topic71 | 484 | 160 | 165 | 809 |
|  | Topic72 | 372 | 130 | 144 | 646 |
|  | Topic73 | 284 | 99 | 114 | 497 |
|  | Topic74 | 398 | 146 | 113 | 657 |
|  | Topic75 | 837 | 277 | 271 | 1385 |
|  | Topic76 | 846 | 278 | 306 | 1430 |
|  | Topic77 | 31 | 13 | 8 | 52 |
|  | Topic78 | 108 | 34 | 31 | 173 |
|  | Topic79 | 620 | 206 | 185 | 1011 |
|  | Topic80 | 766 | 246 | 243 | 1255 |
|  | Topic81 | 702 | 218 | 204 | 1124 |
|  | Topic82 | 411 | 149 | 138 | 698 |
|  | Topic83 | 713 | 230 | 241 | 1184 |
|  | Topic84 | 355 | 111 | 115 | 581 |
|  | Topic85 | 258 | 99 | 84 | 441 |
|  | Topic86 | 283 | 107 | 86 | 476 |
|  | Topic87 | 282 | 85 | 102 | 469 |
|  | Topic88 | 640 | 223 | 206 | 1069 |
|  | Topic91 | 566 | 176 | 156 | 898 |
|  | Topic92 | 661 | 224 | 234 | 1119 |
|  | Topic93 | 539 | 162 | 147 | 848 |
|  | Topic94 | 487 | 144 | 152 | 783 |
|  | Topic95 | 383 | 123 | 115 | 621 |
|  | Topic96 | 333 | 102 | 102 | 537 |
|  | Topic97 | 571 | 190 | 190 | 951 |
|  | Topic98 | 663 | 211 | 231 | 1105 |
|  | Topic99 | 623 | 236 | 256 | 1115 |
|  | Topic100 | 394 | 146 | 139 | 679 |

#### Supplementary Table 2. Summary of performance metrics of binary models

BOM, DNABERT (fine-tuned), LS-GKM, Enformer (fine-tuned) models trained to predict mouse E8.25 distal CREs ^3^. Accuracy, F1 score, auPR, precision, recall and auROC were calculated across each of the 17 cell types in the dataset.

| **Cell type** | **Model** | **Accuracy** | **F1** | **auPR** | **Precision** | **Recall** | **auROC** | **MCC** |
| --- | --- | --- | --- | --- | --- | --- | --- | --- |
| **Allantois** | **BOM** | **0.936** | **0.940** | **0.995** | **0.897** | **0.987** | **0.995** | **0.905** |
|  | DNABERT | 0.516 | 0.590 | 0.527 | 0.519 | 0.684 | 0.519 | 0.027 |
|  | LS-GKM | 0.781 | 0.773 | 0.842 | 0.817 | 0.734 | 0.835 | 0.565 |
|  | Enformer | 0.748 | 0.761 | 0.837 | 0.738 | 0.785 | 0.829 | 0.497 |
| **Cardiomyocyte** | **BOM** | **0.960** | **0.960** | **0.994** | **0.971** | **0.949** | **0.994** | **0.969** |
|  | DNABERT | 0.790 | 0.796 | 0.841 | 0.780 | 0.811 | 0.851 | 0.579 |
|  | LS-GKM | 0.798 | 0.799 | 0.875 | 0.804 | 0.794 | 0.869 | 0.597 |
|  | Enformer | 0.795 | 0.791 | 0.890 | 0.817 | 0.766 | 0.889 | 0.592 |
| **Endothelium** | **BOM** | **0.909** | **0.911** | **0.977** | **0.916** | **0.906** | **0.972** | **0.982** |
|  | DNABERT | 0.826 | 0.829 | 0.851 | 0.831 | 0.828 | 0.879 | 0.651 |
|  | LS-GKM | 0.858 | 0.854 | 0.938 | 0.902 | 0.812 | 0.932 | 0.721 |
|  | Enformer | 0.942 | 0.943 | 0.979 | 0.947 | 0.939 | 0.979 | 0.884 |
| **Erythroid** | **BOM** | **0.928** | **0.930** | **0.977** | **0.915** | **0.946** | **0.980** | **0.971** |
|  | DNABERT | 0.759 | 0.772 | 0.767 | 0.742 | 0.804 | 0.816 | 0.519 |
|  | LS-GKM | 0.779 | 0.782 | 0.836 | 0.781 | 0.784 | 0.853 | 0.557 |
|  | Enformer | 0.918 | 0.920 | 0.978 | 0.909 | 0.931 | 0.979 | 0.836 |
| **ExE endoderm** | **BOM** | **0.887** | **0.862** | **0.942** | **0.862** | **0.862** | **0.959** | **0.890** |
|  | DNABERT | 0.535 | 0.459 | 0.479 | 0.438 | 0.483 | 0.584 | 0.054 |
|  | LS-GKM | 0.578 | 0.651 | 0.790 | 0.491 | 0.966 | 0.845 | 0.340 |
|  | Enformer | 0.859 | 0.853 | 0.961 | 0.744 | 1.000 | 0.977 | 0.753 |
| **Forebrain** | **BOM** | **0.964** | **0.966** | **0.998** | **0.990** | **0.942** | **0.997** | **0.933** |
|  | DNABERT | 0.601 | 0.621 | 0.728 | 0.636 | 0.606 | 0.669 | 0.201 |
|  | LS-GKM | 0.720 | 0.733 | 0.811 | 0.755 | 0.712 | 0.805 | 0.441 |
|  | Enformer | 0.819 | 0.847 | 0.875 | 0.776 | 0.933 | 0.882 | 0.645 |
| **Gut** | **BOM** | **0.945** | **0.948** | **0.992** | **0.948** | **0.948** | **0.990** | **0.948** |
|  | DNABERT | 0.722 | 0.758 | 0.829 | 0.703 | 0.822 | 0.809 | 0.443 |
|  | LS-GKM | 0.769 | 0.765 | 0.876 | 0.828 | 0.711 | 0.867 | 0.546 |
|  | Enformer | 0.851 | 0.866 | 0.951 | 0.826 | 0.911 | 0.945 | 0.703 |
| **Mesenchyme** | **BOM** | **0.910** | **0.909** | **0.963** | **0.914** | **0.905** | **0.965** | **0.934** |
|  | DNABERT | 0.764 | 0.757 | 0.818 | 0.772 | 0.743 | 0.839 | 0.528 |
|  | LS-GKM | 0.778 | 0.771 | 0.854 | 0.790 | 0.752 | 0.856 | 0.557 |
|  | Enformer | 0.859 | 0.859 | 0.924 | 0.851 | 0.867 | 0.93 | 0.717 |
| **Mid/hindbrain** | **BOM** | **0.938** | **0.941** | **0.978** | **0.941** | **0.941** | **0.979** | **0.900** |
|  | DNABERT | 0.627 | 0.651 | 0.625 | 0.644 | 0.659 | 0.638 | 0.251 |
|  | LS-GKM | 0.727 | 0.725 | 0.809 | 0.773 | 0.682 | 0.820 | 0.459 |
|  | Enformer | 0.814 | 0.833 | 0.888 | 0.790 | 0.882 | 0.889 | 0.629 |
| **Mixed mesoderm** | **BOM** | **0.845** | **0.824** | **0.941** | **0.955** | **0.724** | **0.954** | **0.717** |
|  | DNABERT | 0.500 | 0.408 | 0.476 | 0.500 | 0.345 | 0.484 | 0.000 |
|  | LS-GKM | 0.741 | 0.706 | 0.842 | 0.818 | 0.621 | 0.816 | 0.498 |
|  | Enformer | 0.810 | 0.807 | 0.881 | 0.821 | 0.793 | 0.889 | 0.621 |
| **Neural crest** | **BOM** | **0.961** | **0.962** | **0.995** | **0.962** | **0.962** | **0.993** | **0.940** |
|  | DNABERT | 0.612 | 0.615 | 0.665 | 0.637 | 0.594 | 0.639 | 0.225 |
|  | LS-GKM | 0.765 | 0.766 | 0.824 | 0.797 | 0.737 | 0.832 | 0.532 |
|  | Enformer | 0.824 | 0.844 | 0.921 | 0.782 | 0.917 | 0.914 | 0.655 |
| **NMP** | **BOM** | **0.961** | **0.962** | **0.995** | **0.962** | **0.962** | **0.994** | **0.959** |
|  | DNABERT | 0.773 | 0.783 | 0.848 | 0.774 | 0.793 | 0.860 | 0.544 |
|  | LS-GKM | 0.785 | 0.778 | 0.883 | 0.837 | 0.726 | 0.869 | 0.577 |
|  | Enformer | 0.775 | 0.779 | 0.858 | 0.794 | 0.764 | 0.859 | 0.551 |
| **Paraxial mesoderm** | **BOM** | **0.924** | **0.927** | **0.989** | **0.930** | **0.925** | **0.988** | **0.966** |
|  | DNABERT | 0.684 | 0.702 | 0.711 | 0.700 | 0.704 | 0.735 | 0.367 |
|  | LS-GKM | 0.731 | 0.736 | 0.777 | 0.764 | 0.711 | 0.783 | 0.464 |
|  | Enformer | 0.761 | 0.763 | 0.843 | 0.800 | 0.730 | 0.836 | 0.525 |
| **Pharyngeal mesoderm** | **BOM** | **0.882** | **0.884** | **0.963** | **0.871** | **0.897** | **0.966** | **0.854** |
|  | DNABERT | 0.474 | 0.432 | 0.505 | 0.474 | 0.397 | 0.472 | -0.051 |
|  | LS-GKM | 0.719 | 0.725 | 0.790 | 0.714 | 0.735 | 0.790 | 0.437 |
|  | Enformer | 0.756 | 0.759 | 0.802 | 0.754 | 0.765 | 0.812 | 0.511 |
| **Somatic mesoderm** | **BOM** | **0.960** | **0.962** | **0.996** | **0.973** | **0.951** | **0.996** | **0.985** |
|  | DNABERT | 0.472 | 0.000 | 0.566 | 0.000 | 0.000 | 0.548 | 0.000 |
|  | LS-GKM | 0.815 | 0.815 | 0.909 | 0.867 | 0.768 | 0.887 | 0.637 |
|  | Enformer | 0.741 | 0.756 | 0.842 | 0.752 | 0.761 | 0.820 | 0.480 |
| **Spinal cord** | **BOM** | **0.949** | **0.952** | **0.991** | **0.921** | **0.987** | **0.991** | **0.978** |
|  | DNABERT | 0.645 | 0.662 | 0.755 | 0.648 | 0.677 | 0.735 | 0.289 |
|  | LS-GKM | 0.740 | 0.741 | 0.821 | 0.757 | 0.727 | 0.813 | 0.480 |
|  | Enformer | 0.751 | 0.796 | 0.860 | 0.687 | 0.946 | 0.862 | 0.540 |
| **Surface ectoderm** | **BOM** | **0.877** | **0.877** | **0.917** | **0.899** | **0.856** | **0.931** | **0.931** |
|  | DNABERT | 0.713 | 0.720 | 0.821 | 0.720 | 0.720 | 0.791 | 0.426 |
|  | LS-GKM | 0.734 | 0.721 | 0.849 | 0.778 | 0.672 | 0.823 | 0.473 |
|  | Enformer | 0.840 | 0.833 | 0.930 | 0.898 | 0.776 | 0.909 | 0.688 |

#### Supplementary Table 3. Summary of prediction statistics for random mouse E8.25 data splits

Binary BOM models was trained to predict enhancers specific to 17 mouse E8.25 cell types ^3^. For every cell type, five random data splits were produced, dividing at random enhancers into training, validation and test sets. The mean accuracy, F1, auPR, precision, recall and auROC values were calculated across the 17 cell types for every random data split.

| Random data split | Accuracy | F1 | auPR | Precision | Recall | auROC | MCC |
| --- | --- | --- | --- | --- | --- | --- | --- |
| #1 | 0.929 | 0.925 | 0.974 | 0.921 | 0.933 | 0.976 | 0.860 |
| #2 | 0.924 | 0.917 | 0.974 | 0.937 | 0.909 | 0.975 | 0.851 |
| #3 | 0.919 | 0.914 | 0.965 | 0.914 | 0.915 | 0.970 | 0.837 |
| #4 | 0.918 | 0.914 | 0.973 | 0.912 | 0.920 | 0.977 | 0.839 |
| #5 | 0.932 | 0.934 | 0.984 | 0.931 | 0.939 | 0.983 | 0.865 |

#### Supplementary Table 4. Summary of BOM multiclass test results using different error functions

The mean value of each performance metric across E8.25 17 cell types is shown ^3^.

| **eval_metric** | **Recall** | **Precision** | **Accuracy** | **F1** | **auROC** | **auPR** | **MCC** |
| --- | --- | --- | --- | --- | --- | --- | --- |
| mlogloss | 0.882 | 0.985 | 0.995 | 0.926 | 0.999 | 0.988 | 0.927 |
| merror | 0.848 | 0.986 | 0.993 | 0.906 | 0.999 | 0.990 | 0.908 |

#### Supplementary Table 5. Prediction statistics for mouse E8.25 topics

A multiclass BOM model was trained to predict the topic of topic-specific mouse E8.25 CREs ^3^. Mean values of accuracy, F1 score, auPR, precision, recall and auROC were calculated for the predictions of CREs specific to 93 topics.

| Accuracy | F1 | auPR | Precision | Recall | auROC | MCC |
| --- | --- | --- | --- | --- | --- | --- |
| 0.995 | 0.712 | 0.859 | 0.943 | 0.588 | 0.995 | 0.737 |

#### Supplementary Table 6. Summary of performance of DNN multiclass models on test data

| **CNNx3 + FCx2** (Basset) training/fine-tuning | Model Size (number of parameters) | Hyperparameter  selection | | Performance on test set | | | Training log | | |
| --- | --- | --- | --- | --- | --- | --- | --- | --- | --- |
|  |  | lr | batch_size | AVE_auc | AVE_auPR | MCC | Wall_Time | CPU_Time | Epoch |
| CNNx3 + FCx2_without grid search | 3,993,317 | 0.002 | 128 | 0.625 | 0.108 | 0.1132 | 12m50s | 2h0m50s | 29 |
| CNNx3 + FCx2_optimal hyperparameters | 3,993,317 | 0.0026 | 128 | 0.654 | 0.124 | 0.145 | 12m44s | 1h56m50s | 28 |
| **Basset_fine-tuned** | 17,017 | 0.0080 | 64 | **0.807** | **0.279** | **0.322** | 5m33s | 46m48s | 48 |

| **CNN + LSTM** (DeepMel**)** training/fine-tuning | Model Size (number of parameters) | Hyperparameter  selection | | Performance on test set | | | Training log | | |
| --- | --- | --- | --- | --- | --- | --- | --- | --- | --- |
|  |  | lr | batch_size | AVE_auc | AVE_auPR | MCC | Wall_Time | CPU_Time | Epoch |
| **CNN + LSTM_without grid search** | 3,440,401 | 0.001 | 128 | **0.802** | **0.262** | **0.337** | 7m32s | 42m42s | 30 |
| CNN + LSTM_optimal hyperparameter | 3,440,401 | 0.0012 | 128 | 0.789 | 0.245 | 0.320 | 7m36s | 47m41s | 32 |
| DeepMel_fine-tuned | 4,369 | 0.0084 | 64 | 0.660 | 0.132 | 0.098 | 15m39s | 1h36m38s | 48 |

| **CNNx4 + FCx2** training/fine-tuning | Model Size (number of parameters) | Hyperparameter  selection | | Performance on test set | | | Training log | | |
| --- | --- | --- | --- | --- | --- | --- | --- | --- | --- |
|  |  | lr | batch_size | AVE_auc | AVE_auPR | MCC | Wall_Time | CPU_Time | Epoch |
| CNNx4 + FCx2_without grid search | 1,120,113 | 0.002 | 128 | 0.706 | 0.154 | 0.168 | 3m10s | 22m50s | 11 |
| **CNNx4 + FCx2_optimal hyperparameter** | 1,120,113 | 0.0083 | 64 | **0.792** | **0.234** | **0.283** | 3m10s | 22m48s | 11 |

All tasks were executed on a compute node with 12 CPUs and the best performance is highlighted in bold. *_without Grid Search: Model was trained using pre-selected hyperparameters, without performing a grid search. *_with Grid Search: Model was trained using the optimal hyperparameters identified through a grid search. *_fine-tuned: The model was fine-tuned on a pre-trained model, utilizing pre-trained weights and training only the final output layer for specificity.

#### Supplementary Table 7. Summary of performance metrics of mouse E8.25 multiclass models

BOM, Basset (fine-tuned), CNN + LSTM architecture and CNNx4 + FCx2 architecture models trained to predict the cell type of mouse E8.25 enhancers. Accuracy, F1 score, auPR, precision, recall and auROC were calculated for each of the 17 cell types. ‘NA’ means the statistic could not be calculated because of zero denominator.

| Cell type | Model | Accuracy | F1 | auPR | Precision | Recall | auROC | MCC |
| --- | --- | --- | --- | --- | --- | --- | --- | --- |
| Allantois | **BOM** | **0.994** | **0.903** | **0.984** | **1.000** | **0.823** | **0.999** | **0.904** |
|  | Basset | 0.968 | nan | 0.108 | nan | 0.000 | 0.735 | 0.000 |
|  | CNN + LSTM | 0.968 | nan | 0.104 | nan | 0.000 | 0.764 | 0.000 |
|  | CNNx4 + FCx2 | 0.968 | nan | 0.121 | 0.000 | 0.000 | 0.778 | -0.004 |
| Cardiomyocyte | **BOM** | **0.996** | **0.971** | **0.993** | **0.994** | **0.949** | **0.999** | **0.969** |
|  | Basset | 0.930 | 0.065 | 0.221 | 0.545 | 0.034 | 0.784 | 0.124 |
|  | CNN + LSTM | 0.932 | 0.115 | 0.312 | 0.647 | 0.063 | 0.838 | 0.187 |
|  | CNNx4 + FCx2 | 0.928 | 0.032 | 0.222 | 0.300 | 0.017 | 0.798 | 0.057 |
| Endothelium | **BOM** | **0.996** | **0.985** | **0.998** | **0.983** | **0.986** | **1.000** | **0.982** |
|  | Basset | 0.941 | 0.786 | 0.858 | 0.828 | 0.748 | 0.954 | 0.753 |
|  | CNN + LSTM | 0.923 | 0.658 | 0.813 | 0.920 | 0.512 | 0.942 | 0.653 |
|  | CNNx4 + FCx2 | 0.879 | 0.667 | 0.793 | 0.555 | 0.834 | 0.936 | 0.615 |
| Erythroid | **BOM** | **0.996** | **0.973** | **0.996** | **0.966** | **0.980** | **1.000** | **0.971** |
|  | Basset | 0.919 | 0.205 | 0.352 | 0.520 | 0.127 | 0.840 | 0.229 |
|  | CNN + LSTM | 0.924 | 0.195 | 0.524 | 0.719 | 0.113 | 0.903 | 0.265 |
|  | CNNx4 + FCx2 | 0.923 | 0.409 | 0.442 | 0.555 | 0.324 | 0.880 | 0.386 |
| ExE endoderm | **BOM** | **0.998** | **0.885** | **0.972** | **1.000** | **0.793** | **1.000** | **0.889** |
|  | Basset | 0.988 | nan | 0.144 | nan | 0.000 | 0.888 | 0.000 |
|  | CNN + LSTM | 0.988 | nan | 0.062 | nan | 0.000 | 0.789 | 0.000 |
|  | CNNx4 + FCx2 | 0.988 | nan | 0.040 | nan | 0.000 | 0.749 | 0.000 |
| Forebrain | **BOM** | **0.995** | **0.934** | **0.995** | **0.989** | **0.885** | **1.000** | **0.933** |
|  | Basset | 0.960 | 0.108 | 0.249 | 0.857 | 0.058 | 0.813 | 0.216 |
|  | CNN + LSTM | 0.958 | nan | 0.123 | nan | 0.000 | 0.764 | 0.000 |
|  | CNNx4 + FCx2 | 0.958 | nan | 0.095 | nan | 0.000 | 0.738 | 0.000 |
| Gut | **BOM** | **0.995** | **0.951** | **0.993** | **0.969** | **0.933** | **1.000** | **0.948** |
|  | Basset | 0.953 | 0.396 | 0.515 | 0.667 | 0.281 | 0.915 | 0.414 |
|  | CNN + LSTM | 0.946 | nan | 0.123 | nan | 0.000 | 0.729 | 0.000 |
|  | CNNx4 + FCx2 | 0.946 | nan | 0.103 | nan | 0.000 | 0.682 | 0.000 |
| Mesenchyme | **BOM** | **0.995** | **0.935** | **0.981** | **0.979** | **0.895** | **0.999** | **0.934** |
|  | Basset | 0.961 | 0.284 | 0.389 | 0.655 | 0.181 | 0.877 | 0.331 |
|  | CNN + LSTM | 0.957 | nan | 0.106 | 0.000 | 0.000 | 0.753 | -0.004 |
|  | CNNx4 + FCx2 | 0.958 | nan | 0.191 | nan | 0.000 | 0.820 | 0.000 |
| Mid/hindbrain | **BOM** | **0.994** | **0.896** | **0.983** | **1.000** | **0.812** | **0.999** | **0.898** |
|  | Basset | 0.967 | 0.046 | 0.157 | 1.000 | 0.024 | 0.775 | 0.151 |
|  | CNN + LSTM | 0.966 | nan | 0.088 | nan | 0.000 | 0.758 | 0.000 |
|  | CNNx4 + FCx2 | 0.966 | nan | 0.093 | nan | 0.000 | 0.768 | 0.000 |
| Mixed mesoderm | **BOM** | **0.994** | **0.682** | **0.937** | **1.000** | **0.517** | **0.999** | **0.717** |
|  | Basset | 0.988 | nan | 0.033 | nan | 0.000 | 0.613 | 0.000 |
|  | CNN + LSTM | 0.988 | nan | 0.023 | nan | 0.000 | 0.738 | 0.000 |
|  | CNNx4 + FCx2 | 0.988 | nan | 0.033 | nan | 0.000 | 0.741 | 0.000 |
| Neural crest | **BOM** | 0.994 | 0.943 | 0.992 | 0.961 | 0.925 | **1.000** | 0.940 |
|  | Basset | 0.947 | nan | 0.171 | nan | 0.000 | 0.772 | 0.000 |
|  | CNN + LSTM | 0.947 | nan | 0.142 | nan | 0.000 | 0.756 | 0.000 |
|  | CNNx4 + FCx2 | 0.928 | 0.304 | 0.246 | 0.315 | 0.293 | 0.838 | 0.266 |
| NMP | **BOM** | **0.994** | **0.962** | **0.994** | **0.958** | **0.967** | **1.000** | **0.959** |
|  | Basset | 0.916 | 0.119 | 0.271 | 0.583 | 0.066 | 0.776 | 0.176 |
|  | CNN + LSTM | 0.927 | 0.445 | 0.526 | 0.629 | 0.344 | 0.882 | 0.431 |
|  | CNNx4 + FCx2 | 0.924 | 0.264 | 0.428 | 0.739 | 0.160 | 0.851 | 0.322 |
| Paraxial mesoderm | **BOM** | **0.996** | **0.968** | **0.997** | **0.975** | **0.962** | **1.000** | **0.966** |
|  | Basset | 0.939 | 0.105 | 0.284 | 0.750 | 0.057 | 0.816 | 0.195 |
|  | CNN + LSTM | 0.936 | nan | 0.160 | nan | 0.000 | 0.743 | 0.000 |
|  | CNNx4 + FCx2 | 0.936 | nan | 0.132 | nan | 0.000 | 0.733 | 0.000 |
| Pharyngeal mesoderm | **BOM** | **0.993** | **0.848** | **0.982** | **1.000** | **0.735** | **0.999** | **0.854** |
|  | Basset | 0.973 | nan | 0.090 | nan | 0.000 | 0.733 | 0.000 |
|  | CNN + LSTM | 0.973 | nan | 0.106 | nan | 0.000 | 0.766 | 0.000 |
|  | CNNx4 + FCx2 | 0.973 | nan | 0.059 | nan | 0.000 | 0.723 | 0.000 |
| Somitic mesoderm | **BOM** | **0.997** | **0.987** | **0.998** | **0.992** | **0.981** | **1.000** | **0.985** |
|  | Basset | 0.891 | 0.111 | 0.281 | 0.405 | 0.065 | 0.775 | 0.128 |
|  | CNN + LSTM | 0.922 | 0.560 | 0.661 | 0.689 | 0.471 | 0.919 | 0.530 |
|  | CNNx4 + FCx2 | 0.902 | 0.229 | 0.415 | 0.692 | 0.137 | 0.820 | 0.279 |
| Spinal cord | **BOM** | **0.996** | **0.980** | **0.999** | **0.982** | **0.978** | **1.000** | **0.978** |
|  | Basset | 0.910 | 0.082 | 0.256 | 0.500 | 0.045 | 0.781 | 0.129 |
|  | CNN + LSTM | 0.917 | 0.290 | 0.392 | 0.627 | 0.188 | 0.848 | 0.313 |
|  | CNNx4 + FCx2 | 0.912 | 0.180 | 0.377 | 0.558 | 0.108 | 0.836 | 0.218 |
| Surface ectoderm | **BOM** | **0.994** | **0.932** | **0.995** | **0.991** | **0.88** | **1.000** | **0.931** |
|  | Basset | 0.951 | 0.281 | 0.366 | 0.522 | 0.192 | 0.869 | 0.296 |
|  | CNN + LSTM | 0.950 | nan | 0.183 | nan | 0.000 | 0.743 | 0.000 |
|  | CNNx4 + FCx2 | 0.950 | 0.074 | 0.192 | 0.500 | 0.040 | 0.780 | 0.131 |

#### Supplementary Table 8. Summary of prediction statistics of mouse E8.5 enhancers using models trained on mouse E8.25 enhancers

Mouse E8.5 enhancers specific to 15 cell types were scored using binary BOM models trained on mouse E8.25 enhancers to distinguish similar cell types. The models were trained on enhancers trimmed to their central 500bp and E8.5 enhancers were trimmed in a similar way. Mean values of accuracy, F1, auPR, precision, recall and auROC values were calculated across the predictions produced by the 15 models.

| Accuracy | F1 | auPR | Precision | Recall | auROC | MCC |
| --- | --- | --- | --- | --- | --- | --- |
| 0.759 | 0.692 | 0.847 | 0.831 | 0.606 | 0.852 | 0.532 |

#### Supplementary Table 9. Summary of prediction statistics for different motif detection thresholds

Binary BOM models were trained to distinguish enhancers specific to 17 mouse E8.25 cell types and 6 human cell lines. We trained the models on datasets produced with different motif detection thresholds (q-value <= 0.1, q-value <= 0.3 and q-value <= 0.5). Mean values of accuracy, F1 score, auPR precision, recall and auROC were calculated for each dataset and every motif detection threshold (N = 17 and 6 mouse E8.25 cell types and human cell lines, respectively).

| Dataset | q-value threshold | Accuracy | F1 | auPR | Precision | Recall | auROC | MCC |
| --- | --- | --- | --- | --- | --- | --- | --- | --- |
| Mouse E8.25 | <= 0.1 | 0.882 | 0.883 | 0.956 | 0.891 | 0.880 | 0.952 | 0.766 |
|  | <= 0.3 | 0.936 | 0.937 | 0.983 | 0.939 | 0.936 | 0.982 | 0.873 |
|  | <= 0.5 | 0.934 | 0.933 | 0.984 | 0.933 | 0.937 | 0.984 | 0.869 |
| Human cell lines | <= 0.1 | 0.688 | 0.670 | 0.774 | 0.707 | 0.666 | 0.769 | 0.392 |
|  | <= 0.3 | 0.869 | 0.872 | 0.948 | 0.848 | 0.902 | 0.950 | 0.743 |
|  | <= 0.5 | 0.923 | 0.926 | 0.978 | 0.893 | 0.962 | 0.980 | 0.850 |

Supplementary Table 10. Summary of prediction statistics of BOM models trained on overlapping and non-overlapping TF binding motifs counts

Binary BOM models were trained to predict enhancers specific to each of the 17 mouse E8.25 cell types. Mean accuracy, F1 score, auPR, precision, recall and auROC values were calculated across the 17 models.

| Model motifs | Accuracy | F1 | auPR | Precision | Recall | auROC | MCC |
| --- | --- | --- | --- | --- | --- | --- | --- |
| Non-overlapping | 0.602 | 0.532 | 0.686 | 0.680 | 0.454 | 0.664 | 0.235 |
| overlapping | 0.934 | 0.933 | 0.984 | 0.933 | 0.937 | 0.984 | 0.869 |

Supplementary Table 11. Prediction statistics for human cell lines

Binary BOM models were trained to predict cell line specific CREs against a background composed of CREs from the other cell lines. We used a total of 6 cell lines (Gm12878, Hela-S3, Huvec, H1-hESC, HepG2 and K562). We calculated accuracy, F1 score, auPR, precision, recall and auROC across the 6 models. We used q-value <= 0.5 as motif detection threshold.

| Accuracy | F1 | auPR | Precision | Recall | auROC | MCC |
| --- | --- | --- | --- | --- | --- | --- |
| 0.923 | 0.926 | 0.978 | 0.893 | 0.962 | 0.980 | 0.850 |

#### Supplemental Table 12. Summary of prediction statistics of human hematopoiesis enhancers

Binary BOM models were trained to predict enhancers specific to 22 cell types. Mean accuracy, F1 score, auPR, precision, recall and auROC values were calculated across the 22 data sets.

| Accuracy | F1 | auPR | Precision | Recall | auROC | MCC |
| --- | --- | --- | --- | --- | --- | --- |
| 0.897 | 0.901 | 0.949 | 0.873 | 0.934 | 0.958 | 0.799 |

#### Supplemental Table 13. Summary of prediction statistics of zebrafish enhancers

Binary BOM models were trained to predict enhancers specific to 11 adult zebrafish tissues. Mean values of accuracy, F1, auPR, precision, recall and auROC across the 11 test sets are shown.

| Accuracy | F1 | auPR | Precision | Recall | auROC | MCC |
| --- | --- | --- | --- | --- | --- | --- |
| 0.960 | 0.961 | 0.991 | 0.941 | 0.983 | 0.992 | 0.921 |

#### Supplementary Table 14. Human fetal enhancers prediction statistics

Binary BOM models were trained to predict human fetal enhancers specific to cardiomyocytes and erythroblast cells. The values of accuracy, F1 score, auPR, precision, recall and auROC are shown for each model.

| Cell type | Accuracy | F1 | auPR | Precision | Recall | auROC | MCC |
| --- | --- | --- | --- | --- | --- | --- | --- |
| Cardiomyocytes | 0.949 | 0.952 | 0.974 | 0.915 | 0.992 | 0.981 | 0.901 |
| Erythroblasts | 0.873 | 0.851 | 0.931 | 0.952 | 0.769 | 0.923 | 0.755 |

#### Supplementary Table 15. Cross-species prediction statistics of cell type specific CREs

Binary BOM models were trained to distinguish human or mouse cell-type specific CREs cardiomyocyte (‘CM’) and erythroblasts (‘ER’). The models were tested in CREs from a similar cell type of the other species. The mean accuracy, F1 score, auPR, precision, recall and auROC values are shown in each case.

| Species (model) | Species (test data) | Cell type | Accur. | F1 | auPR | Prec. | Recall | auROC | MCC |
| --- | --- | --- | --- | --- | --- | --- | --- | --- | --- |
| Mouse | Human | CM | 0.755 | 0.723 | 0.835 | 0.821 | 0.645 | 0.850 | 0.520 |
|  |  | ER | 0.706 | 0.688 | 0.845 | 0.759 | 0.629 | 0.813 | 0.421 |
| Human | Mouse | CM | 0.711 | 0.652 | 0.833 | 0.823 | 0.540 | 0.814 | 0.450 |
|  |  | ER | 0.763 | 0.743 | 0.805 | 0.809 | 0.688 | 0.817 | 0.532 |

#### Supplementary Table 16. Summary of prediction performance metrics for BOM models trained using different tree depth values

We used the mouse E8.25 cell type-specific enhancers dataset to evaluate the effect of the maximum decision tree depth in the classification performance (XGBoost parameter “max_depth”). BOM binary models were trained for each of the 17 cell types using a maximum tree depth of 6 (default), 8, 10 and 12. The mean values of accuracy, F1 score, auPR, precision, recall and auROC across the 17 cell types are shown.

| Decision trees depth | Accuracy | F1 | auPR | Precision | Recall | auROC | MCC |
| --- | --- | --- | --- | --- | --- | --- | --- |
| 6 (default) | 0.934 | 0.933 | 0.984 | 0.933 | 0.937 | 0.984 | 0.869 |
| 8 | 0.941 | 0.939 | 0.986 | 0.940 | 0.942 | 0.986 | 0.882 |
| 10 | 0.938 | 0.937 | 0.986 | 0.933 | 0.945 | 0.986 | 0.877 |
| 12 | 0.938 | 0.937 | 0.985 | 0.935 | 0.943 | 0.985 | 0.877 |
